# An NR2F1-dependent retinoic acid network controls retinal specialization in mice and reveals foveal hypoplasia in patients with BBSOAS

**DOI:** 10.64898/2026.09.22.753406

**Authors:** Paolo Piovani, Carole Belliardo, Rahul Makam, Boglarka Zambo, Annabelle Mantilleri, Amandine Chassot, Gergo Gogl, Christian P. Schaaf, Seth Blackshaw, Patrick Yu-Wai-Man, Michele Bertacchi, Michèle Studer

**Author notes:** Co-last and co-corresponding authors.

## Abstract

The molecular programs that establish specialized retinal regions during development are essential for high-acuity vision, yet how their disruption contributes to human visual disorders remains poorly understood. Bosch-Boonstra-Schaaf optic atrophy syndrome (BBSOAS), caused by pathogenic variants in *NR2F1* and characterized by visual impairment, provides an opportunity to investigate these mechanisms. Using single-cell RNA sequencing of three complementary *Nr2f1* mouse models, including two carrying patient-specific mutations, we identified a shared *Nr2f1*-dependent transcriptional program enriched in retinoic acid (RA) pathway genes. Loss or mutation of *Nr2f1* disrupted the spatial organization of RA signaling, most prominently by expanding the dorso-equatorial *Cyp26a1* expression domain into ventral retina and reducing ventral determinants such as *Vax2*. These molecular changes were associated with altered dorso-ventral distribution of S-and M-opsin-expressing cone photoreceptors. We further demonstrate that human NR2F1 binds a conserved regulatory region upstream of *CYP26A1*, supporting its direct role in regulating local RA availability. Finally, high-resolution optical coherence tomography in individuals with BBSOAS revealed reproducible foveal abnormalities, including a smaller and shallower foveal pit and increased central retinal thickness, consistent with foveal hypoplasia. These findings uncover a previously unrecognized retina-intrinsic component of BBSOAS visual pathology and establish an NR2F1-RA/CYP26A1 regulatory axis linking developmental retinal regionalization to human foveal specialization.

## INTRODUCTION

The developmental programs that generate specialized retinal territories required for high-acuity vision remain poorly understood, particularly in the context of human neurodevelopmental disorders (NDDs) (*1*). Through coordinated regulation of regional identity, progenitor competence, and neuronal differentiation, retinal patterning establishes distinct functional domains that ultimately support color vision, spatial resolution, and central retinal specialization, including formation of the fovea in primates (*2–4*). Identifying the molecular signals and gene regulatory networks controlling these processes is therefore critical for understanding how developmental perturbations lead to visual impairment.

In mice, retinal neurogenesis begins around embryonic day (E) 11.5 and follows a highly conserved developmental sequence that generates the six neuronal classes and Müller glia, reaching completion during the second postnatal week (*5*, *6*). This intrinsic program is progressively refined by patterning cues acting along the dorso-ventral (D-V) and naso-temporal (N-T) retinal axes (*2*). Among the signaling pathways governing retinal regionalization, BMP (*7*), WNT (*8*), SHH (*9*), and retinoic acid (RA) (*10*, *11*) have emerged as key morphogenetic regulators. RA signaling plays a central role in optic cup morphogenesis (*10*, *12*), ventral retinal identity (*13*), retinal progenitor competence, and neuronal subtype specification (*12–14*). Its disruption is associated with human ocular malformations (*15–18*). Unlike classical morphogens, RA activity is regulated through discrete domains defined by the coordinated expression of RA-synthesizing aldehyde dehydrogenase (ALDHs) and RA-degrading CYP26 enzymes. This tightly controlled distribution of RA activity is essential for specifying retinal regional identity. Beyond its role in early patterning, RA also contributes to cone photoreceptor subtype specification and central-peripheral retinal specialization (*19*), processes essential for high-acuity vision. Remarkably, the presumptive primate fovea coincides with a localized suppression of RA signaling mediated by restricted CYP26 expression in foveal progenitors, suggesting that precise spatial control of RA activity is a prerequisite for foveal development (*20*). Despite these observations, the transcriptional mechanisms that govern the regional distribution of RA activity during retinal development, and whether their disruption contributes to impaired foveal specification, remain largely unknown.

Nuclear receptors represent a major class of transcriptional regulators that integrate extracellular signaling pathways with developmental gene expression programs. Among them, NR2F1 is an evolutionarily conserved member of this family with established roles in eye and cortical development (*21–24*). During early eye development, *Nr2f1* is expressed in the neural retina, optic stalk, and optic nerve, where it contributes to retinal ganglion cell differentiation and survival, and optic nerve development (*25*, *26*). Consistent with these functions, *Nr2f1* mouse models display multiple visual-system abnormalities, including defects in optic vesicle patterning, retinal ganglion cells, optic nerve development, visual acuity, and higher-order visual centers, recapitulating several ocular features of BBSOAS (*35*, *42*). However, these models have not defined the molecular mechanisms by which *NR2F1* regulates retinal regionalization or revealed how disease-associated mutations alter its function. Importantly, patient-specific *Nr2f1* mouse models have only recently become available (*27*, *28*), and their impact on retinal development remains unexplored. Thus, the *Nr2f1*-dependent transcriptional networks that establish retinal regional identity and specialization during early retinogenesis remain largely unknown.

At the molecular level, NR2F1 modulates RA-dependent transcription through interactions with RAR/RXR complexes, binding to RA response elements (RAREs) and recruiting transcriptional corepressors to fine-tune RA-responsive gene expression in a context-and stage-dependent manner (*29*, *30*). NR2F1 also shares overlapping DNA-binding specificity with its homolog NR2F2. However, whether this regulatory activity operates during retinal development remains poorly understood. In particular, it is unknown whether NR2F1 contributes to the spatial control of RA signaling required for retinal regional identity and whether disruption of this regulatory function contributes to the visual abnormalities associated with *NR2F1* haploinsufficiency.

The relevance of NR2F1-dependent developmental mechanisms to human vision is highlighted by Bosch-Boonstra-Schaaf Optic Atrophy Syndrome (BBSOAS) (*31–33*), an NDD caused by NR2F1 haploinsufficiency resulting from whole-gene deletions, truncating variants, or missense mutations affecting either the ligand-binding domain (LBD), required for dimerization and co-factor interactions, or the DNA-binding domain (DBD), which contains two highly conserved zinc fingers essential for DNA recognition (*34*). In addition to hypotonia, autism spectrum disorder, epilepsy, global developmental delay, and intellectual disability (*34*, *35*), visual abnormalities are among the most consistent and debilitating features of BBSOAS (*26*, *31–38*). Affected individuals display reduced visual acuity (*26*, *33*), optic nerve abnormalities including atrophy or hypoplasia (*25*, *39*), refractive errors (*31–33*), and cortical visual impairment (*40*). Although visual deficits are relatively stable (*26*) and may partially improve with therapy (*35*), they substantially impact quality of life. Despite the growing number of reported patients and pathogenic *NR2F1* variants (*26*, *31–33*, *41*, *42*), it remains unknown whether altered retinal specialization contributes to visual dysfunction in BBSOAS or how NR2F1-dependent developmental programs shape retinal organization.

Here, we combine single-cell transcriptomics of embryonic retinae from three complementary *Nr2f1* mouse models with structural retinal analyses of BBSOAS patients, to define the developmental programs controlled by *Nr2f1/NR2F1*. We identify an *Nr2f1*-dependent transcriptional network that spatially regulates RA signaling through control of *Cyp26a1* and other RA pathway genes, thereby shaping retinal regional identity and cone photoreceptor patterning. Importantly, our findings position *NR2F1* as a key upstream regulator of retinal specialization and reveal previously unrecognized foveal hypoplasia in patients with BBSOAS, providing a mechanistic link between disrupted embryonic retinal development and impaired human visual function.

## RESULTS

### Patient-specific mouse models show altered levels of Nr2f1 retinal expression

Individuals with BBSOAS carry heterogenous *NR2F1* deletions or loss-of-function point mutations falling in distinct protein domains, impairing its activity as a transcriptional regulator (*31–34*). To investigate how distinct types of mouse *Nr2f1* genetic alterations impact retinal development, we compared three mouse lines (*28*) (**Fig. 1A,B**): (i) a constitutive knock-out (*Nr2f1 Null*) line in which exon3 and the poly-A signal are deleted (*25*, *41*); (ii) a point mutation in the LBD leading to premature protein truncation (in mouse *E397*\*; in human *E400\**), and (iii) a missense mutation in the DBD causing a single amino-acid substitution (in mouse *R139L;* in human *R142L*) (*28*). For each line, heterozygous (*HET*) or homozygous knock-out/knock-in (*HOM KO/KI*) embryos were generated and systematically compared with line-specific wild-type (*WT*) littermates to minimize variability due to genetic background.

**Fig. 1.**
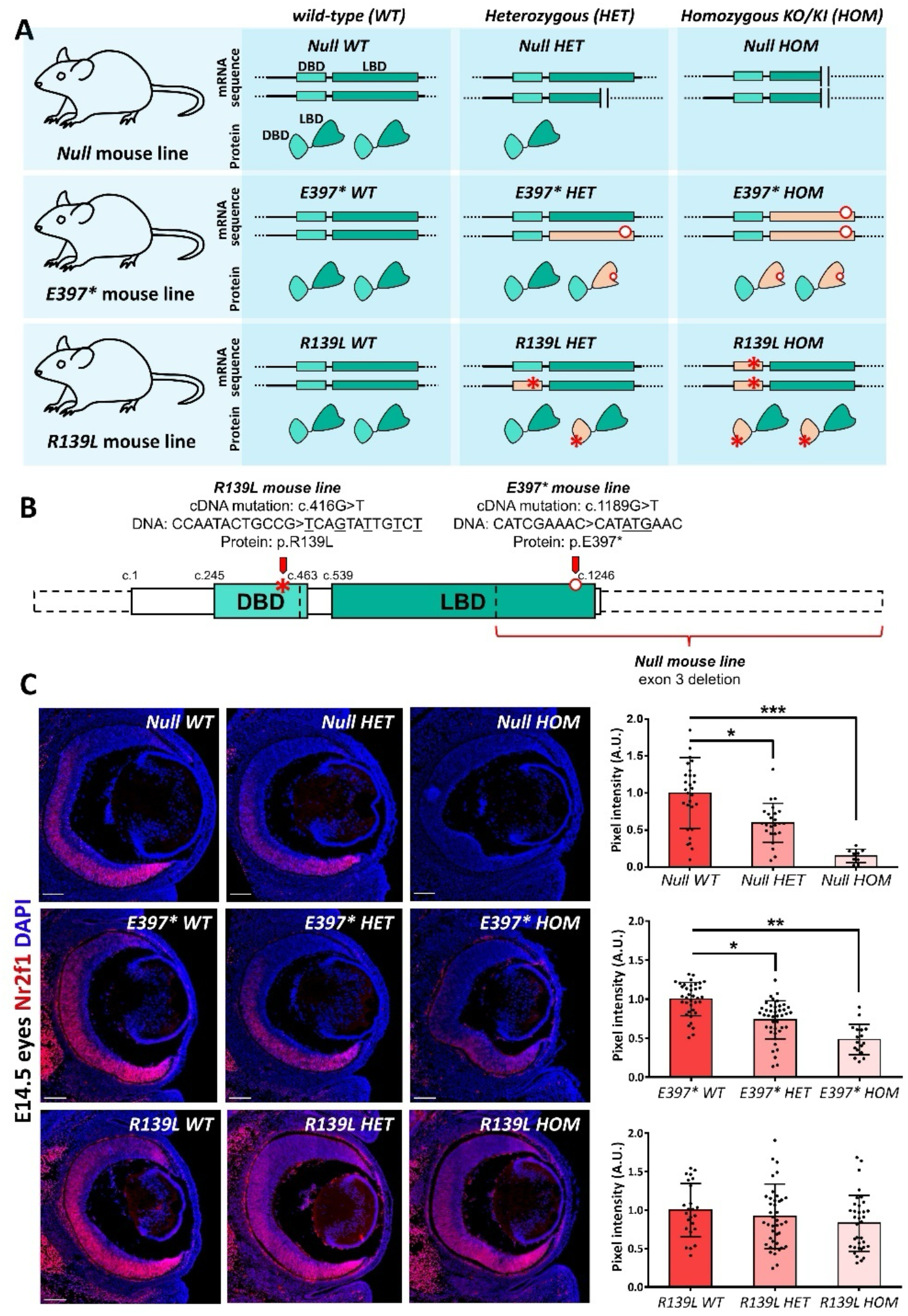
Generation and characterization of the three *Nr2f1* mouse models. **A)** Three *knock-out* (*KO*) or *knock-in* (*KI*) mouse models (*Nr2f1 Null, Nr2f1 E397*, and Nr2f1 R139L*) were used to generate *WT, HET, and HOM* embryos, modelling reduced, absent, or mutated *Nr2f1* function *in vivo.* For each mouse line, schematics illustrate the genotype of the two alleles and the predicted amount of protein produced. **B)** Schematic of the Nr2f1 protein showing the genetic modifications introduced in the three mouse lines and their localization. The protein structure is represented by solid lines superimposed on dotted lines indicating the corresponding mRNA exons. **C)** Immunofluorescence shows Nr2f1 expression (red) in E14.5 eyes. Nr2f1 was undetectable in *Null HOM* embryos and reduced in *Null HET*, E397* *HET* and *E397* HOM* eyes. *R139L HOM* eyes showed a non-significant trend toward reduced Nr2f1 protein levels, to approximately 80% of *WT* levels. Quantification of Nr2f1 pixel intensity is shown in the graphs on the right. Scale bars: 100 µm. Points in graphs correspond to pixel intensity measurements in individual sections; n = 11 to 44 sections from 6 to 8 eyes per genotype. Statistical significance was determined by one-way ANOVA with Tukey’s correction; * p < 0.05; ** p < 0.01; *** p < 0.001.

First, *Nr2f1* mRNA expression was assessed across genotypes using allele-specific primers targeting either the *WT* or mutant alleles (**Fig. S1A**). Primers specific for the *E397\** and *R139L* mutant sequences confirmed expression of the mutant alleles in *HET* and *HOM* embryos (**Fig. S1B,C**), while primers recognizing the corresponding *WT* sequences confirmed reduced *Nr2f1* mRNA levels in these mutants (**Fig. S1D,E**). Similarly, primers targeting the excised exon3 detected reduced transcript levels in *Null HET* and *HOM* tissues (**Fig. S1F**). Notably, primers amplifying a region common to all genotypes detected residual *Nr2f1* mRNA in *Null HOM* embryos, indicating that transcripts lacking exon3 and the polyadenylation signal persist at low levels, although they are likely rapidly degraded (**Fig. S1G**). In contrast, total *Nr2f1* mRNA levels were increased in both *E397\** and *R139L HET* and *HOM* embryos, as detected by the universal primer pair (**Fig. S1G**), suggesting a compensatory transcriptional response to *Nr2f1* inactivation. Because *Nr2f1* mRNA levels differed between mutant lines, we next assessed Nr2f1 protein abundance by quantifying immunofluorescence pixel intensity in embryonic retinae (**Fig. 1C**). Consistent with previous reports (*25*, *26*, *43*), Nr2f1 was strongly expressed throughout the retina, displaying a ventral-high to dorsal-low gradient. As expected, Nr2f1 protein levels were reduced in *Null HET* and undetectable in *Null HOM* retinae (**Fig. 1C**). In contrast, Nr2f1 protein remained detectable in both *E397\** and *R139L HOM* eyes (**Fig. 1C**), indicating that truncated or mutated Nr2f1 proteins were produced *in vivo*, although at reduced levels, particularly in *HOM E397\** retinae. Notably, whereas Nr2f1 normally localizes to the nucleus (*44*), the truncated E397* Nr2f1 protein showed aberrant cytoplasmic localization in retinal cells (**Fig. S1H**), suggesting impaired nuclear targeting *in vivo*, consistent with previous observations in cell culture (*45*).

Collectively, these data reveal genotype-dependent effects of *Nr2f1* mutations on Nr2f1 protein abundance in the embryonic retina, with reduced protein levels despite increased *Nr2f1* mRNA in some mutants, suggesting reduced protein stability and compensatory regulation of *Nr2f1* mRNA.

### Single-cell transcriptomics defines the spatial organization of *Nr2f1*-expressing retinal progenitor cells

Previous studies have reported an altered balance between retinal progenitor cells (RPCs) and post-mitotic retinal ganglion cells (RGCs) in the ventral-most region of the *Nr2f1 Null* retina (*25*), where Nr2f1 is normally expressed at its highest levels during embryonic development. To determine whether truncated or mutant Nr2f1 proteins exert similar effects, we quantified the number of Vsx2+ RPCs and Tuj1+ RGCs in the ventral retina of E14.5 *E397\** and *R139L HET* and *HOM* embryos and compared these with the *Null* line (**Fig. S2A**). While *HET* embryos showed no significant alterations, *HOM* embryos exhibited a pronounced imbalance between mitotic and post-mitotic cells in the ventral retina (**Fig. S2B**), indicative of impaired retinogenesis following *Nr2f1* loss or mutation. We therefore selected E14.5 for the following analyses, as this mid-retinal developmental stage encompasses both proliferating RPCs and newly generated post-mitotic neurons.

Hence, to define the cellular and molecular consequences of Nr2f1 loss, truncation, or DBD mutation at single-cell resolution, we performed scRNA-Seq analysis on E14.5 retinae from *WT*, *HET*, and *HOM* embryos across the three mouse models using 10X Genomics technology. Initial clustering identified retinal and non-retinal populations based on established *bona fide* markers, including retinal cells (*Rax*+, *Vsx2*+, *Six6*+), RGCs (*Tubb3*+, *Pou4f1*+), lens (*Foxe3*+), RPE (*Mitf*+), connective tissue (*Pitx2*+), endothelial cells (*Pecam1*+), epidermal cells (*Krt5*+) and microglia (*Aif1*+) (**Fig. S3A,B**). We then focused on retinal cells and closely associated lens and RPE populations by retina-centered re-clustering (**Fig. 2A,B**), resolving 30 clusters organized around proliferative RPCs (*Pax6*+, *Vsx2*+, *Mki67*+). From this progenitor pool, intermediate neurogenic precursors (*Atoh7*+) branched into three major differentiation trajectories: *Map2*+/*Pou4f1*+ RGCs, *Crx*+ presumptive photoreceptors, and *Prdm13*+/*Ptf1a*+ and *Lhx1*+ presumptive AC/HCs. Notably, *Nr2f1* expression was highest in RPCs and progressively decreased in more differentiated cell types, suggesting a prominent role for *Nr2f1* in early undifferentiated progenitors (**Fig. S3C**). Consistent with our qRT-PCR analyses (**Fig. S1A-G**), *Nr2f1* expression was reduced in *Null HET* and *HOM* samples, whereas *E397\** and *R139L HET* and *HOM* RPCs showed increased *Nr2f1* transcripts levels (**Fig. S3C,D**), further supporting a compensatory transcriptional response to mutant *Nr2f1* alleles.

**Fig. 2.**
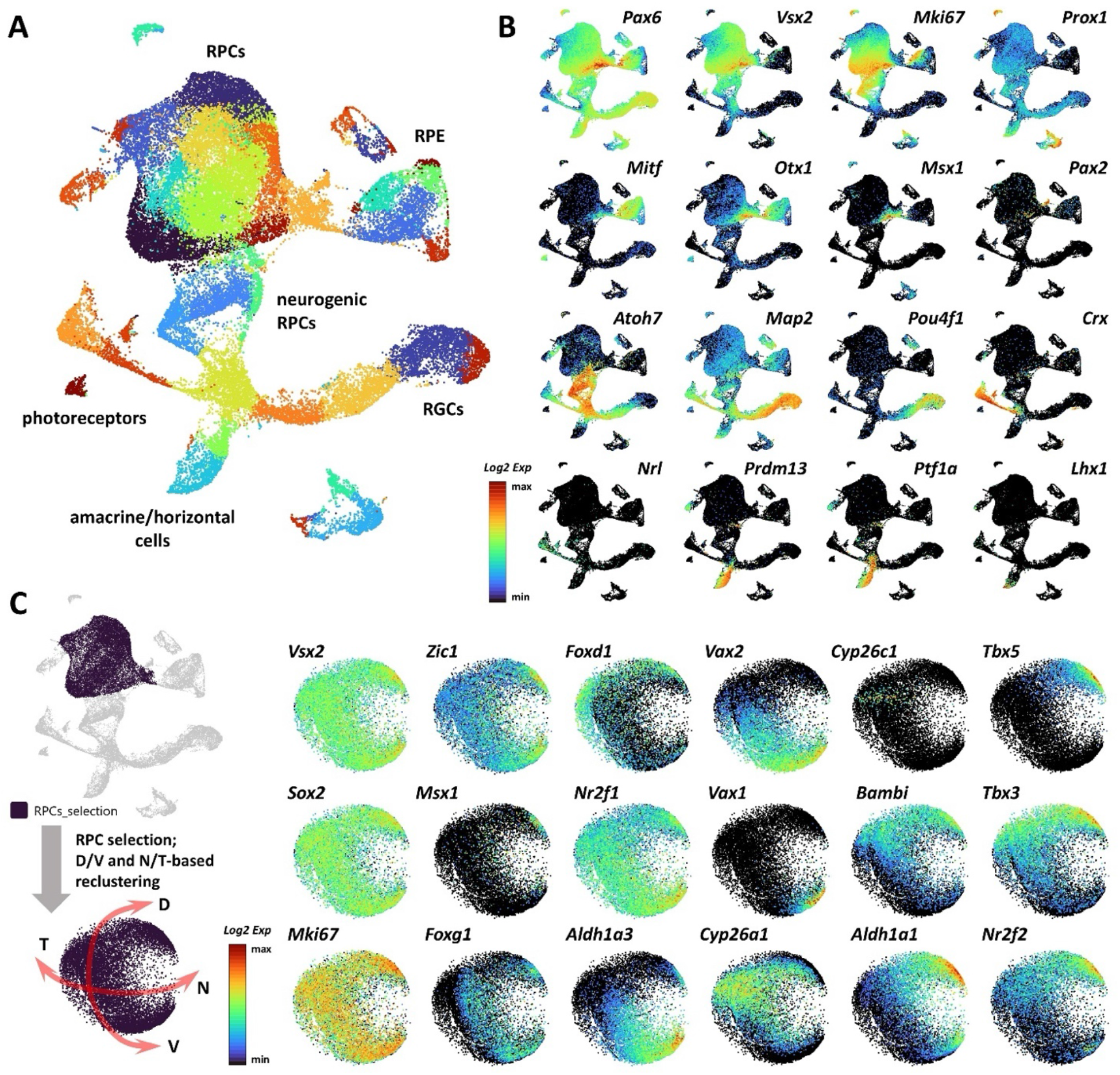
Single-cell transcriptomics defines the spatial organization of *Nr2f1*-expressing retinal progenitor cells (RPCs). A,B) UMAP projection of re-clustered scRNA-Seq data comprising retinal cells, retinal pigment epithelium (RPE), and lens cells. Retinal populations were subdivided into proliferating RPCs (*Pax6*⁺, *Vsx2*⁺, *Mki67*⁺), neurogenic RPCs (*Atoh7*⁺), and post-mitotic differentiating populations, including *Map2*⁺, *Pou4f1*⁺ retinal ganglion cells (RGCs), *Crx*⁺ photoreceptor precursors, and *Prdm13*⁺, *Ptf1a*⁺, and *Lhx1*⁺ amacrine and horizontal cells. **C)** Left, schematic representation of the RPC population selected for re-clustering and pseudo-spatial annotation based on the expression of *bona fide* naso-temporal (N-T) and dorso-ventral (D-V) regional markers in *WT* retinae. Right, selected UMAP projections showing the expression levels of *Sox2* and *Vsx2* (general RPC markers), *Mki67* (proliferating RPCs), *Zic1* and *Msx1* (CMZ progenitors), *Foxg1* (nasal RPCs), *Foxd1* (temporal RPCs), *Nr2f1, Aldh1a3, Vax2*, and *Vax1* (ventral RPCs), *Cyp26a1* and *Cyp26c1* (equato-dorsal RPCs), and *Bambi, Aldh1a1, Tbx5, Tbx3,* and *Nr2f2* (dorsal RPCs), revealing distinct RPC populations with regional identities that recapitulate the spatial organization of the undifferentiated retina.

To specifically examine the molecular consequences of *Nr2f1* loss or mutation in RPCs, we selected *bona fide* RPCs based on *Vsx2* and *Sox2* expression (**Fig. 2C**). RPCs were spatially re-oriented along the dorso-ventral (D-V) and naso-temporal (N-T) axes according to the expression of *bona fide* regional identity marker genes defining dorsal (*Bambi*, *Aldh1a1*, *Tbx5*, *Tbx3*, and *Nr2f2*), equato-dorsal (*Cyp26a1* and *Cyp26c1*), ventral (*Nr2f1*, *Aldh1a3*, *Vax2*, and *Vax1*), nasal (*Foxg1*), and temporal (*Foxd1*) domains. This approach generated a coherent pseudo-spatial organization of RPCs that recapitulated the major retinal axes. Proliferating Mki67⁺ RPCs were distributed throughout the retinal configuration, whereas Zic1⁺ and Msx1⁺ populations were more restricted to ciliary margin zone (CMZ)-like clusters (**Fig. 2C**).

Together, these results establish a high-resolution, single cell-level landscape of embryonic retinal neurogenesis, spanning proliferating RPCs, neurogenic precursors, and differentiating retinal neuronal lineages. They further identify RPCs as the major cellular compartment expressing *Nr2f1* and provide a pseudo-spatial framework along the D-V and N-T axes for investigating how Nr2f1 disruption alters the transcriptional programs underlying retinal regional identity.

### *Nr2f1* variants differentially alter retinal gene expression depending on mutation type

Differential gene expression analysis was performed by comparing *WT* RPCs with their mutant counterparts. Because patients with BBSOAS are heterozygous for pathogenic *NR2F1* variants (*46*), we initially focused on *HET* animals, comparing each *HET* mutant with its corresponding line-specific *WT* controls (*WT versus HET*) (**Fig. S4**). Strikingly, the *R139L* line exhibited several hundred unique differentially expressed genes (DEGs), far exceeding the number identified in the other two models (**Fig. S4A**). This suggests that the DBD *R139L* mutation exerts a particularly strong impact on the *Nr2f1* downstream transcriptional program than either the E397* truncation or *Null* allele loss, consistent with previous observations linking *R139L* to the most severe anatomical and behavioral phenotypes in mice (*28*).

To distinguish mutation-specific from shared transcriptional alterations, DEG lists from the three mouse lines were intersected using Venn diagram analysis (**Fig. S4B,C**). Only five DEGs were shared across all three lines, and included *Nr2f1* itself, *Prss23*, *Lgr5*, *Glo1* and *Crabp2* (**Fig. S4D,E**), revealing a limited core transcriptional signature associated with heterozygous *Nr2f1* disruption. The large set of *R139L*-specific DEGs was enriched for gene ontology categories related to nervous system development, including axon guidance, cell migration and regulation of apoptosis (**Fig. S4F,G**). Together, these findings demonstrate that the transcriptional consequences of *Nr2f1* disruption are strongly mutation-dependent, with the DBD *R139L* variant producing a markedly broader transcriptional response than either the *E397\** truncation or simple allele loss in the *Null* model. Consistent with previous reports (*28*), these results further support a genotype-phenotype correlation in which specific missense mutations can exert effects that extend beyond those expected from simple reduction of *Nr2f1* dosage.

### Nr2f1 loss or mutation disrupts a shared transcriptional network controlling D-V and N-T retinal patterning

Heterozygous (*HET*) murine models could display milder phenotypes than humans carrying the corresponding heterozygous *NR2F1* variants, reflecting species-specific differences in gene dosage sensitivity and developmental robustness, as well as potential compensatory mechanisms. We therefore reasoned that homozygous disruption or mutation of *Nr2f1* would produce more pronounced molecular phenotypes and facilitate DEG identification and compared control *WT* RPCs with those from *Null HOM* embryos lacking *Nr2f1* or *E397\** and *R139L HOM* embryos carrying two mutant alleles (*WT versus HOM* DEGs). Hundreds of up-or downregulated DEGs were identified in each line (**Fig. 3A**; **Fig. S5A**), with differences in the proportion of RPCs expressing individual DEGs (**Fig. S5B**). Separate GO analysis revealed both shared and line-specific functional categories (**Fig. S5C**). *Null HOM* mutants were enriched for categories related to “dysregulation of apoptotic processes” and “generation of neurons” categories, whereas *E397\** mutants showed broader categories related to “brain development”. As observed in their *HET* counterparts, *R139L HOM* mutants were enriched for “cell migration” and “axon guidance” processes. These findings indicate that different *Nr2f1* mutations differentially affect developmental transcriptional programs.

**Fig. 3.**
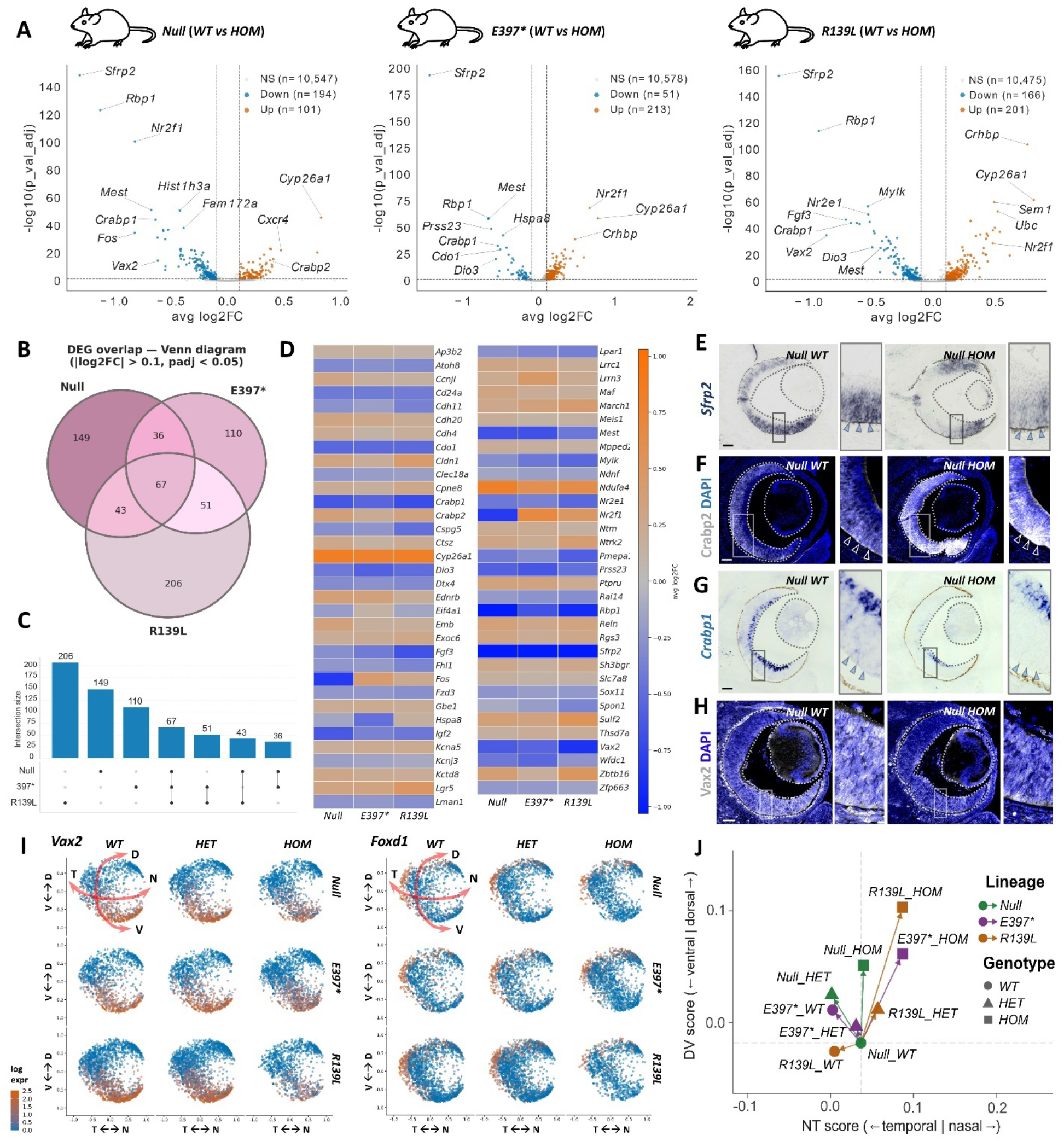
*Nr2f1* loss or mutation disrupts a shared transcriptional network controlling retinal regional identity. **A)** Volcano plots showing differentially expressed genes (DEGs) identified by comparing *WT* and *HOM KO/KI* RPCs in the three mouse models, as indicated, with fold change (log₂FC) on the x-axis and statistical significance (−log₁₀ adjusted p value) on the y-axis. **B,C)** Venn diagram (B) and intersection scheme (C) illustrating the overlap among DEGs identified in the three mouse models. Sixty-seven DEGs are shared in the inner core of the Venn diagram across models, representing transcripts consistently dysregulated in mutant compared with control retinae. **D)** Expression changes of the 67 shared DEGs, with color coding indicating fold change (log_2_FC). **E–H)** *In situ* hybridizations and immunostainings for Sfrp2 (E), Crabp2 (F), Crabp1 (G), and Vax2 (H) on E14.5 eye sections from *Null WT* (left) and *Null HOM KO* (right) embryos. Arrowheads in high-magnification insets indicate regions showing more pronounced increases or reductions in expression. Complete analyses of *Sfrp2, Crabp1, Crabp2,* and *Vax2* across additional genotypes and N-T levels are presented in *Fig. S6.* **I)** Expression levels of the ventral determinant *Vax2* and the temporal determinant *Foxd1* in spatially re-oriented RPCs from the indicated genotypes. Reduced *Vax2* and *Foxd1* expression in mutant retinae indicates partial loss of ventral and temporal patterning identity, respectively. **J)** RPC regional identity visualized according to D-V (y-axis) and N-T (x-axis) patterning scores, calculated from the expression of *Foxg1* (nasal RPCs), *Foxd1* (temporal RPCs), *Nr2f1*, *Aldh1a3*, *Vax2*, and *Vax1* (ventral RPCs), *Cyp26a1* and *Cyp26c1* (equato-dorsal RPCs), and *Bambi*, *Aldh1a1*, *Tbx5*, *Tbx3*, and *Nr2f2* (dorsal RPCs). Scores were centered on *Null WT* retinae as the reference point. Scale bars: 50 µm.

To distinguish mutation-specific from shared transcriptional changes, the identified DEGs were organized in a Venn diagram (**Fig. 3B,C**). We reasoned that the 67 DEGs shared across all three models could represent a core *Nr2f1*-responsive transcriptional program that is disrupted irrespective of the mutation type. Most shared DEGs showed concordant up-or downregulated across the three models, indicating that *Nr2f1* loss or mutation similarly affects their expression (**Fig. 3A,D**). A notable exception was *Nr2f1* itself, which was increased in *E397\** and *R139L HOM* retinae but reduced in *Null HOM* eyes (**Fig. 3D**), as observed by RT-PCR **(Fig. S1**). Several shared DEGs were key regulators of early RPC identity, including *Mest*, *Dio3*, and *Fgf3*, consistent with a role for *Nr2f1* in regulating early-stage temporal identity in RPCs (*47*) (**Fig. 3D**). These early-stage retinal progenitor-enriched genes also included *Sfrp2* as one of its most strongly downregulated transcripts (*48*). *Sfrp2* was reduced in *Null HOM* retinae and showed an even stronger decrease in *E397\** and *R139L HET* and *HOM* retinae (**Figs. 3D,E** and **Fig. S6A**). *In situ* hybridization (ISH) on mouse eye sections showed that *Sfrp2* transcript levels were particularly reduced in central and ventral retinal regions and were completely abolished in the ventral retina of *R139L HOM* embryos, consistent with the stronger molecular phenotype associated with the DBD mutation (**Fig. S6A**). ISH and immunostaining further confirmed dysregulation of highly modulated genes such as *Crabp1* and *Crabp2* (**Fig. 3F,G** and **Fig. S6B,C**), with *Crabp2* showing strong upregulation on the ventral side of the most nasal regions of mutant retinae (**Fig. S6C**). Notably, we also identified marked alterations in the expression of the ventral identity gene *Vax2*, which has an established role in D-V retinal patterning and whose disruption causes retinal developmental abnormalities (*49*, *50*), (**Fig. 3H**; **Fig. S6D**). Both ISH and immunostaining confirmed reduced Vax2 expression and contraction of its ventral domain (**Fig. S6D**), suggesting abnormal D-V patterning and partial loss of ventral retinal identity.

Given the altered expression of D-V determinants and the differential modulation of regional genes along the D-V and N-T axes, we next re-organized the RPC clusters according to their regional identities to visualize key D-V and N-T factors in control and mutant retinae (**Fig. 3I**) and developed a patterning score to quantitively assess regional identity across samples (**Fig. 3J**). Mutant retinae, particularly those from *HOM* animals, separated from their corresponding controls, reflecting a shift in RPC identity toward dorsal characteristics. In the case of *E397\** and *R139L HOM* retinae, this dorsalization was accompanied by a shift toward nasal identity at the expense of temporal identity (**Fig. 3J**).

Together, these data define an *Nr2f1*-dependent transcriptional program core including key regulators of retinal development and regional identity. Disruption of this network occurs independently of the type of *Nr2f1* perturbation and is associated with altered RPC temporal competence and aberrant regional identity along both the D-V and N-T axes.

### *Nr2f1* loss-or mutation converges on dysregulation of the retinoic acid (RA) pathway

To further define the molecular nature of the shared *Nr2f1*-dependent retinal network, we performed gene ontology analysis on the 67 shared core DEGs across the three mouse models. In addition to expected biological process categories such as “generation of neurons” and “nervous system development”, this analysis identified “retinoid binding” as a significantly enriched molecular function (**Fig. S7A**). This finding was of particular interest given the established role of retinoids and RA pathway genes in retinal patterning and cell fate specification along the D-V axis (*51*). Consistently, over-representation analysis (ORA), which assesses the enrichment of functional categories within a gene set relative to a defined background (*52*), identified significant enrichment of RA pathway genes, including *Crabp1, Crabp2, Cyp26a1*, and *Pdk1*. Additional RA-related genes (*Rbp1, Rai14*) were identified by complementary manual curation to account for potential limitations in pathway annotation and ORA, as reported (*53*) (**Fig. 4A**). Furthermore, extending the ORA analysis to all DEGs, including genes belonging to the lateral Venn diagram subsets, identified additional RA-associated genes, such as *Fabp5* in the *Null HOM*, *Aldh1a3, Pdhb* and *Rara* in *E397* HOM*, and *Aldh1a7* in *R139L HOM* RPCs (**Fig. 4B**), suggesting mutation-dependent alterations in RA pathway components. Several of these genes were altered not only in expression level but also in the proportion of RPCs in which they were detected. Notably, *Cyp26a1*, encoding a cytochrome P450 enzyme that degrades RA and is normally expressed in the equatorial/dorsal retina (*20*, *54*), showed an approximately twofold increase in the proportion of *Cyp26a1*+ RPCs in *HOM KO/KI* mutant retinae (**Fig. 4C**; **Fig. S7B**). When RPCs were spatially re-oriented, this increase was associated with a pronounced expansion of the *Cyp26a1*+ RPC population toward the ventral retina (**Fig. 4D**). *Cyp26c1*, a paralog of *Cyp26a1*, was similarly increased, with its normally restricted expression domain becoming more prominent in equatorial/dorsal retinal regions (**Fig. 4D**). Conversely, the proportion of RPCs expressing the ventral marker *Vax2*, a known regulator of *Cyp26a1*(*55*), was reduced across all lines (**Fig. S7B**), consistent with the decreased Vax2 expression detected by immunostaining (**Fig. S6D**).

**Fig. 4.**
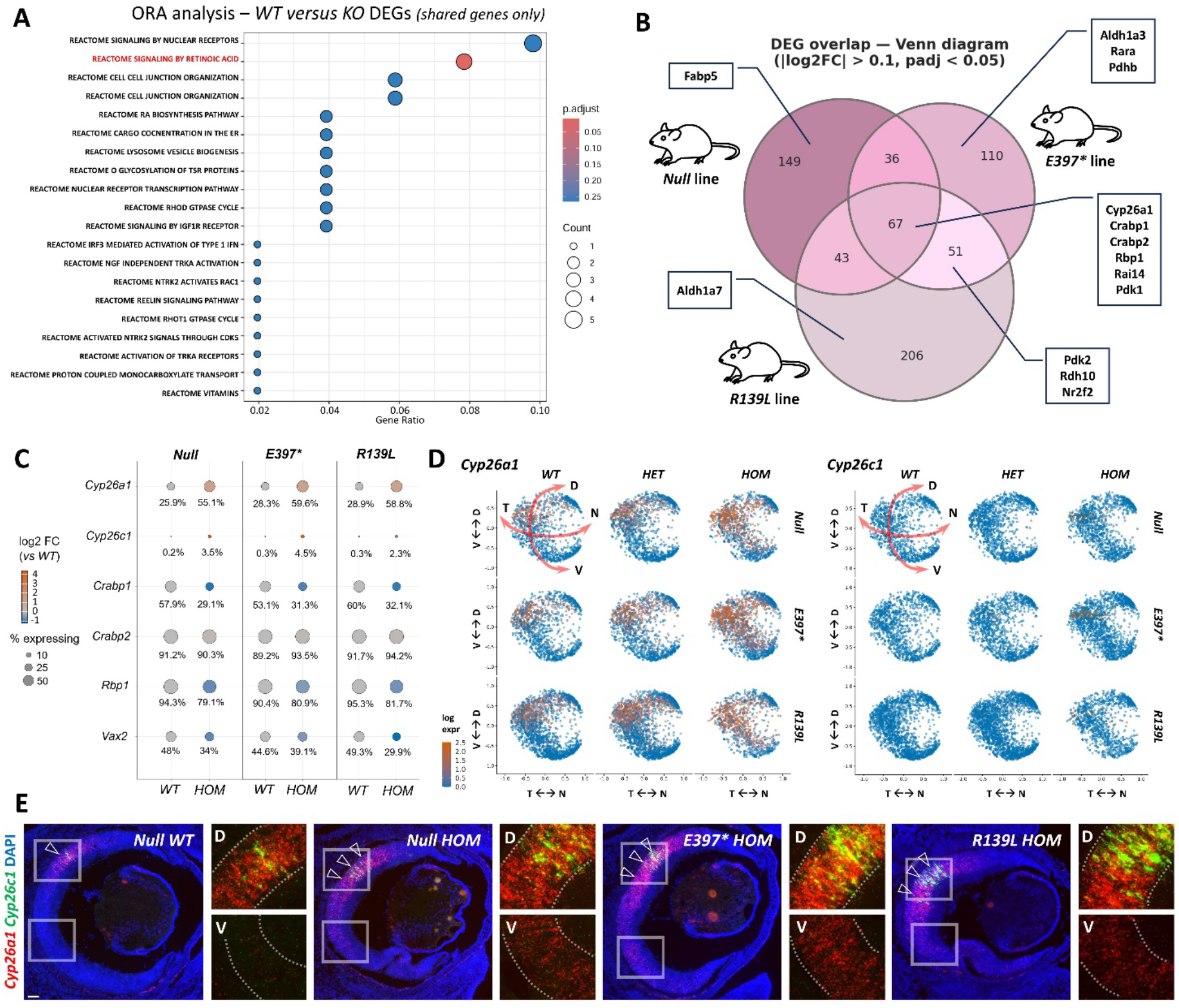
*Nr2f1* disruption alters the spatial regulation of retinoic acid (RA) pathway genes. **A)** Over-representation analysis (ORA) of the 67 *WT versus HOM* DEGs shared across the three mouse models, highlighting significant enrichment of genes associated with the RA pathway. **B)** RA-related genes mapped onto the Venn diagram, including genes dysregulated in only one or two mouse models, revealing both shared and mutation-dependent effects on RA pathway components. **C)** Dot plot showing the proportion of cells expressing the RA-related genes *Cyp26a1, Cyp26c1, Crabp1, Crabp2, Rbp1,* and *Vax2*, together with their fold change in *WT* vs *HOM* RPCs. A complete panel including *HET* retinae is shown in *Fig. S7B.* **D)** Expression levels of the equato-dorsal RA-degrading enzymes *Cyp26a1* and *Cyp26c1* in spatially re-oriented RPCs from the indicated genotypes. Cyp26a1 expression is markedly expanded toward the ventral retina in mutant RPCs. **E)** RNAscope *in situ* hybridization for *Cyp26a1* (red) and *Cyp26c1* (green) in E14.5 retinae from the indicated genotypes. Arrowheads indicate the equato-dorsal expression domain. In addition to upregulation in the dorsal retina of both homologs, ectopic *Cyp26a1* expression is detected in the ventral retina (high-magnification insets; D = dorsal, V = ventral). Scale bars: 50 µm.

Because *Cyp26a1* was the most strongly and significantly upregulated RA-related pathway gene in mutant retinae, both in expression level (**Fig. 3A**) and in the proportion of *Cyp26a1*+ RPCs (**Fig. S7B**), we next examined its spatial expression domain in greater detail. High-sensitivity RNAscope hybridization analysis of control and mutant E14.5 eye sections revealed that, while *Cyp26a1* expression was largely restricted to the equatorial/dorsal domain in *WT* retinae, *HOM* retinae displayed increased expression within this domain, together with a concomitant ventral expansion of the *Cyp26a1*⁺ territory (**Fig. 4E**). *Cyp26c1* expression was also increased in the dorsal retina, with a transition from sparse expression in *WT* to a larger population of expressing cells in mutant retinae (**Fig. 4E**), consistent with the spatially annotated scRNA-Seq data (**Fig. 4D**). Notably, spatial expansion of *Cyp26a1* and *Cyp26c1* expression in the ventral retina, particularly within the ventral CMZ, was most pronounced in temporal retinal regions (**Fig. S7C**). In addition, ectopic *Cyp26a1* expression was detected in the nasal half of the central retina in horizontal sections (**Fig. S7D**). Thus, *Nr2f1* loss of function disrupts the normal spatial restriction of RA-degrading enzymes along both the D-V and N-T axes.

Together, these results highlight dysregulation of the RA pathway as a convergent molecular consequence of distinct *Nr2f1* perturbations. In particular, the marked expansion of the *Cyp26a1* expression domain provides a potential mechanism through which *Nr2f1* disruption could alter local RA availability and thereby impact on retinal regional patterning.

### *Nr2f1* disruption alters M-and S-opsin cone distribution in the post-natal retina

Given the prominent dysregulation of RA pathway genes and the strong upregulation of *Cyp26a1* following *Nr2f1* loss or mutation, we next investigated whether these molecular changes were associated with altered retinal regionalization and cone photoreceptor patterning. Local retinal levels of *Cyp26a1* and other D-V RA pathway components are known to influence cell-type specification and distribution (*55–58*), notably that of cone photoreceptors (*59*). We therefore examined M-opsin and S-opsin distribution by immunostaining retinal sections from *HOM* mutants and their corresponding *WT* controls at different stages of post-natal development, when cone photoreceptors undergo maturation (**Fig. 5**).

**Fig. 5.**
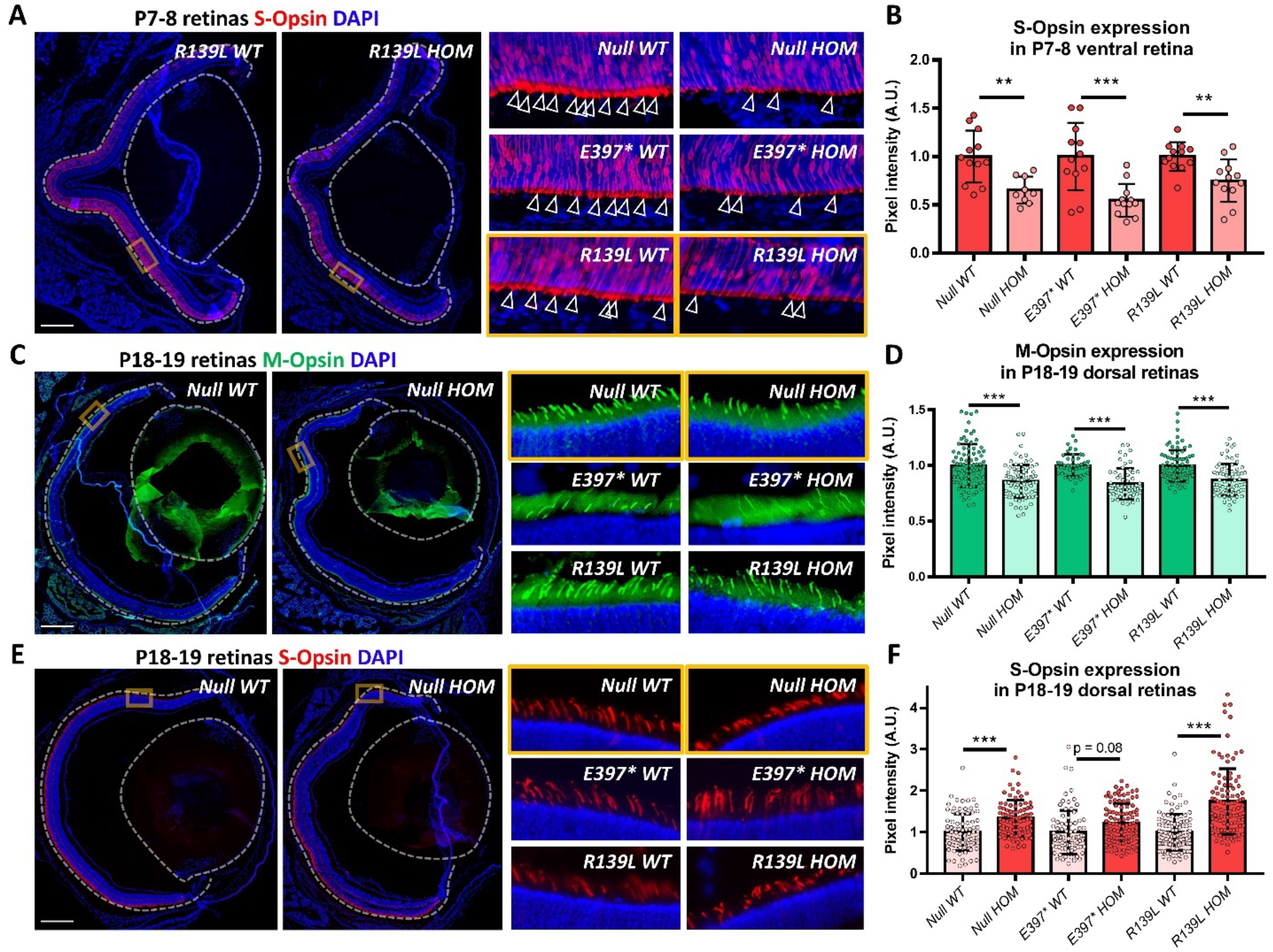
Altered cone opsin expression and D-V distribution in *Nr2f1* mutant retinae. A,B) Immunostaining for short-wavelength opsin (S-Opsin; red) in P7–8 retinae from animals of the indicated genotypes. Representative high-magnification images show reduced S-opsin signal in cone outer segments (arrowheads), with quantification of S-opsin pixel intensity in the ventral retina shown in B. **C,D)** Immunostaining for medium-wavelength opsin (M-opsin; green) in P18-19 retinae. Representative high-magnification images show reduced M-opsin signal in cone outer segments, with quantification of M-opsin pixel intensity in the dorsal retina shown in D. **E,F)** S-opsin immunostaining (red) in P18-19 retinae. Representative images and high-magnification insets show ectopic S-opsin expression in the dorsal-most retina of *Nr2f1* mutants, with quantification of S-opsin pixel intensity in the dorsal retina shown in F. Data are shown for *Null*, *E397*, and *R139L* homozygous (*HOM*) mutants and their corresponding *WT* controls. Scale bars: 500µm. For the graph in B, points correspond to the average pixel intensity measured within 100-µm-wide selections of the outer segment layer; n = 9-12 measurements from n = 3-4 sections from n = 2-4 eyes per genotype. For the graphs in D and F, points correspond to pixel intensity measurements from individual outer segments; n = 80-105 measurements from n = 3-4 sections from n = 2-4 eyes per genotype. Statistical significance was assessed by one-way ANOVA with Tukey’s correction; ** p < 0.01; *** p < 0.001.

At P7-8, when S-opsin expression becomes established in ventral cone photoreceptors, S-opsin immunoreactivity in the ventral retina was significantly reduced in all three *Nr2f1* mutant models compared with their respective *WT* controls (**Fig. 5A,B**). Quantification of fluorescence intensity within cone outer segments revealed reductions of approximately similar magnitude across the three mutant genotypes, indicating that *Nr2f1* disruption consistently impaired S-opsin expression during early post-natal cone maturation. At P18-19, M-opsin expression in the dorsal retina was likewise significantly reduced in all three mutant models compared with their corresponding controls (**Fig. 5C,D**). The reduction was evident both in the immunostaining pattern and in quantitative measurements of M-opsin fluorescence intensity in cone outer segments. Together with the reduction in ventral S-opsin expression observed at P7-8, these findings show that *Nr2f1* disruption affects the expression of both major cone opsins, consistent with its D-V organization of the post-natal mouse retina.

In addition to reduced M-opsin expression, *Nr2f1* disruption altered the normal spatial distribution of S-opsin expression at P18–19 (**Fig. 5E**). Whereas S-opsin expression in *WT* retinae was largely confined to ventral cones, mutant retinae displayed ectopic S-opsin expression extending into the dorsal-most retina. Quantification of dorsal S-opsin fluorescence confirmed a significant increase in *Null HOM* and *R139L HOM* animals compared with their respective controls (**Fig. 5F**), while *E397*\* *HOM* retinae showed a similar trend that did not reach statistical significance (*p* = 0.08). Thus, the consequences of *Nr2f1* disruption extend beyond changes in overall opsin abundance to include altered D-V cone opsin distribution.

Together, these results demonstrate that *Nr2f1* disruption affects both the expression level and spatial distribution of cone opsins during post-natal retinal maturation. In light of the embryonic D-V patterning defects and dysregulation of RA pathway components observed in *Nr2f1* mutant retinae, these findings support a model in which *Nr2f1*-dependent regional patterning during embryogenesis contributes to the subsequent spatial organization of cone subtype identity and opsin expression in the post-natal retina.

### NR2F1 directly binds a highly conserved regulatory region of the *CYP26A1* promoter

The increased *Cyp26a1* expression observed following *Nr2f1* loss or mutation prompted us to investigate whether the two genes occupy complementary expression domains in the *WT* retina. As previously shown, Nr2f1 is expressed during retinal development in a ventral-high to dorsal-low gradient (**Fig. 1**). Consistent with this pattern, RNAscope detection of *Cyp26a1* combined with Nr2f1 immunostaining revealed largely complementary expression domains (**Fig. 6A**), with *Cyp26a1* expression increasing as the Nr2f1 gradient declined toward the dorsal retina. Notably, a similar complementary expression was also observed along the N-T axis. In horizontal retinal sections, Nr2f1 levels were higher in the nasal aspect of the dorsal retina, whereas *Cyp26a1* expression was sparse and predominantly detected in the opposite temporal region, where Nr2f1 levels were lower (**Fig. S8A**). Together, these complementary expression patterns suggested that Nr2f1 may participate in the spatial restriction of *Cyp26a1*. This regulation could occur indirectly, through modulation of *Vax2*, which has been implicated in the control of *Cyp26a1* expression (*55*), and/or through direct regulation of the *Cyp26a1* regulatory region. Thus, *Nr2f1*-dependent spatial expression gradient may contribute to confining *Cyp26a1* expression to its physiological domain (**Fig. 6B**).

**Fig. 6.**
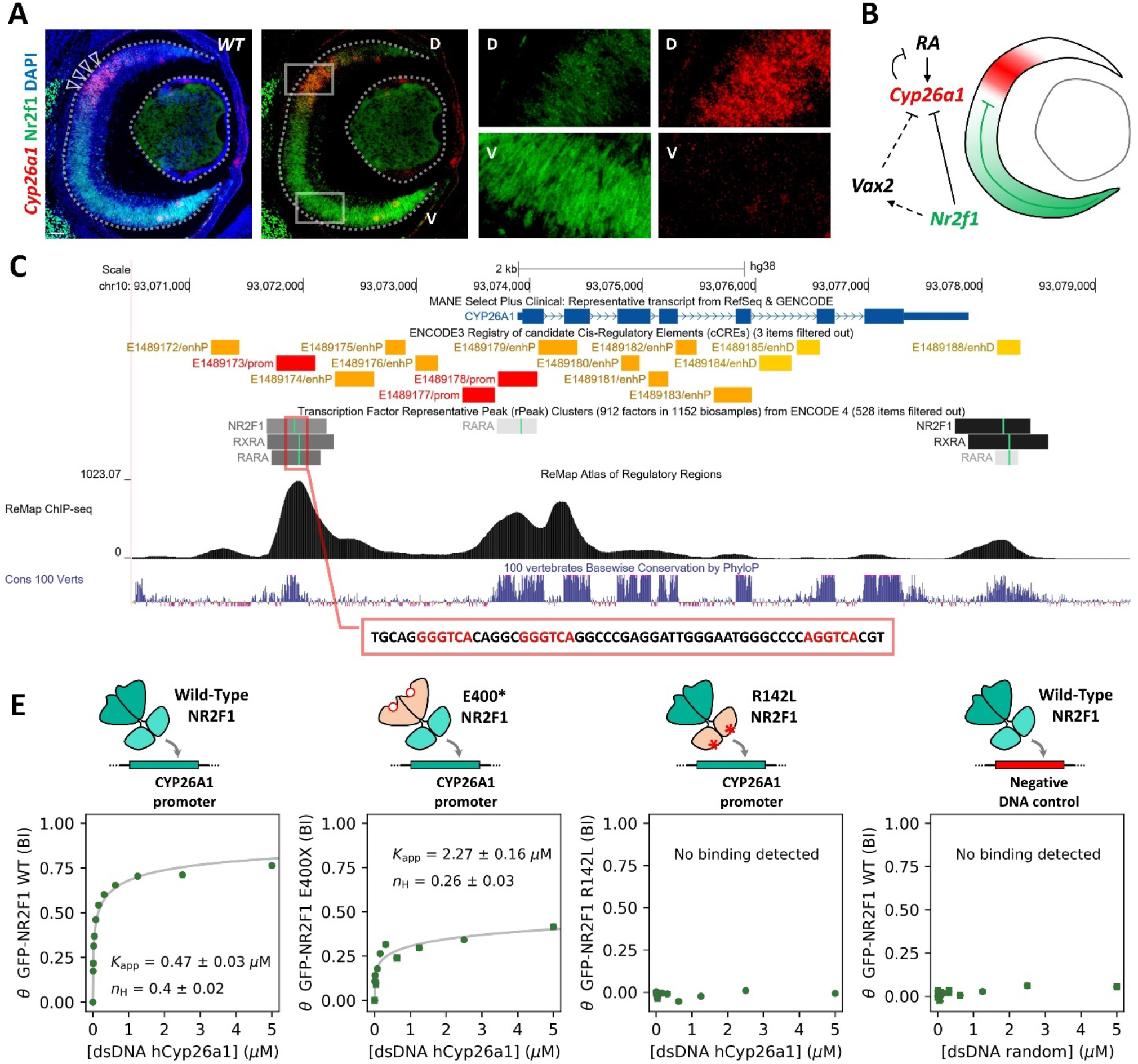
NR2F1 directly binds a regulatory region of *CYP26A1* and may contribute to its spatial restriction. **A)** RNAscope *in situ* hybridization for *Cyp26a1* (red, arrowheads) combined with Nr2f1 immunostaining (green) on E14.5 WT eye sections, showing complementary expression gradients. *Cyp26a1* is expressed in a defined stripe dorsal to Nr2f1, where Nr2f1 expression is lowest (see insets; D = dorsal, V = ventral). **B)** Schematic model of the Nr2f1-dependent regulatory network controlling RA signaling in the mouse retina. Nr2f1 loss or mutation is associated with reduced Vax2 expression and expansion of the Cyp26a1 expression domain toward the ventral retina. **C)** Bioinformatic analysis of the human *CYP26A1* upstream regulatory region (promoter regions; prom; in red), showing putative binding sites for various nuclear receptors, including NR2F1, RARs, and RXRs. Nuclear receptor binding at this region is supported by published ChIP-Seq datasets (ReMap ChIP-Seq peaks), and the region is highly conserved across species (Cons). The 56-bp sequence highlighted in the red box contains the predicted NR2F1 binding sites and was selected for the native holdup assay. **D)** Native holdup assay measuring binding of NR2F1 proteins to the 56-bp *CYP26A1* regulatory sequence. *WT* NR2F1 binds the target sequence, whereas binding is markedly reduced by the E400* LBD truncation and abolished by the R142L DBD mutation. The fitted binding isotherms (grey lines) show a negative cooperative binding model; the reduced Hill coefficient observed with E400* may indicate an altered oligomeric state. An unrelated DNA target bait served as experimental negative control (right). BI, bound fraction intensity; Kapp, apparent affinity; nH, Hill coefficient. Scale bars: 50 µm.

To investigate whether direct DNA binding could underlie this regulation, we searched the human *CYP26A1* promoter region using the UCSC Genome Browser for NR2F1 binding sites. Notably, *in silico*-predicted and ChIP-Seq-validated NR2F1 binding sites were identified within an upstream regulatory region of the *CYP26A1* gene, described as a promoter region (**Fig. 6C**). This evolutionarily conserved region also contained validated RXR and RAR binding sites (**Fig. 6C**), consistent with the established role of RA in regulating *CYP26A1* expression (*60*, *61*).

We next tested whether NR2F1 directly binds this 56-bp-long region using native holdup (*62*, *63*), a sensitive approach for estimating equilibrium binding constants of protein-DNA interactions directly from cell extracts under near-physiological conditions, allowing the capture of genuine protein-DNA interactions. As this technique relies on a GFP-tagged version of the protein of interest (GFP-NR2F1), we first validated that GFP-NR2F1 retained the DNA-binding properties of endogenous NR2F1. To this end, native holdup assays directed against a known NR2F1 target sequence (*64*) were performed in parallel using extracts from NR2F1-expressing neuronal progenitor cells and GFP-NR2F1-transfected HEK293T cells (**Fig. S8B-D**). Similar binding profiles were obtained, confirming that the tagged recombinant protein preserves the binding characteristics of the endogenous protein (**Fig. S8C,D**). We then screened the binding ability of *WT* and mutated NR2F1 proteins to the 56-bp *CYP26A1* promoter sequence bearing NR2F1 putative binding sites. The GFP-tagged *WT* form of human NR2F1 efficiently bound this target sequence (**Fig. 6E**), indicating that the *CYP26A1* regulatory region is directly recognized by NR2F1 in human cells. We next asked whether the patient-associated mutations examined in the mouse models affect this interaction. Native holdup assays were performed using human NR2F1 proteins carrying the corresponding mutations, namely GFP-tagged forms of LBD-mutated NR2F1 E400* and DBD-mutated NR2F1 R142L. The E400* truncation markedly reduced NR2F1 binding to the *CYP26A1* promoter sequence (**Fig. 6E**). More strikingly, the DBD R142L point mutation completely abolished the ability of NR2F1 to bind the *CYP26A1* promoter sequence (**Fig. 6E**).

These data demonstrate that NR2F1 directly binds an evolutionarily conserved regulatory region of the *CYP26A1* promoter region and that disease-associated LBD and DBD mutations differentially impair this interaction. Together with the complementary spatial expression patterns observed *in vivo*, these data support a model in which NR2F1 directly contributes to the spatial restriction of *CYP26A1* expression (**Fig. 6B**). This regulation may involve direct transcriptional control and/or competition with RAR/RXR complexes at neighboring RA response elements, providing a potential molecular mechanism linking *NR2F1* mutations to the ectopic expansion of *Cyp26a1* observed in mutant retinae.

### Foveal hypoplasia is a previously unrecognized feature of BBSOAS and extends beyond canonical optic neuropathy

Unlike mice, whose retina lacks a specialized region for high-acuity vision, most diurnal vertebrates develop a cone-enriched central retinal specialization that supports high-acuity vision and is referred to as the area centralis, visual streak, or fovea, depending on the species (*65*, *66*). Across species, spatially restricted RA signaling and the activity of Cyp26a1 orthologs have been implicated in the sequential regulation of rod versus cone (*67–69*), and cone subtype specification (*67*), thereby contributing to the establishment of central retinal specializations, including the fovea. Given our finding that *Nr2f1* regulates the spatial organization of RA pathway components and cone photoreceptor patterning, we asked whether individuals with BBSOAS might exhibit developmental abnormalities of the fovea. Visual impairment in BBSOAS has mainly been attributed to optic nerve abnormalities or central brain defects up until now (*31*, *34*), and potential abnormalities of the central retina have not been systematically investigated.

To address this, we characterized foveal morphology in seven individuals with BBSOAS using optical coherence tomography (OCT) and compared quantitative foveal morphological parameters with those of eight age-matched controls (**Tables S1, S2**). Remarkably, foveal pit morphology was altered in BBSOAS (**Fig. 7A-G; Table S3**). Central foveal thickness was increased in BBSOAS compared with controls (243.4 ± 23.0 vs 214.2 ± 10.3 µm; *p* = 0.015), consistent with persistence of inner retinal tissue at the foveola. Foveal inner retinal thickness was nearly doubled in BBSOAS (18.9 ± 13.6 vs 10.2 ± 2.1 µm) although this difference did not remain significant after correction for multiple testing (*p* = 0.065). Foveal outer retinal thickness was also increased (223.4 ± 18.4 vs 203.3 ± 9.9 µm; *p* = 0.025). More strikingly, the foveal pit was substantially smaller and shallower in BBSOAS. Pit depth was reduced by 43% (79.9 ± 34.6 vs 139.2 ± 16.1 µm; *p* = 0.003), pit volume by 58% (0.05 ± 0.03 vs 0.12 ± 0.03 mm³; *p* = 0.003), and pit area by 57% (0.03 ± 0.02 vs 0.07 ± 0.01 mm²; *p* = 0.002). The foveal rim was also lower (323.3 ± 19.1 vs 353.3 ± 12.6 µm; *p* = 0.010) and located closer to the foveal center (rim radius 0.89 ± 0.18 vs 1.14 ± 0.14 mm; *p* = 0.015), resulting in a smaller rim disk area (2.60 ± 0.83 vs 4.03 ± 0.97 mm²; *p* = 0.015). Mean pit slope was similarly reduced (5.1 ± 1.9° vs 7.1 ± 1.2°; *p* = 0.039). Together, these measurements reveal a smaller, shallower foveal pit associated with increased central retinal thickness in individual with BBSOAS, consistent with abnormal foveal development.

**Fig. 7.**
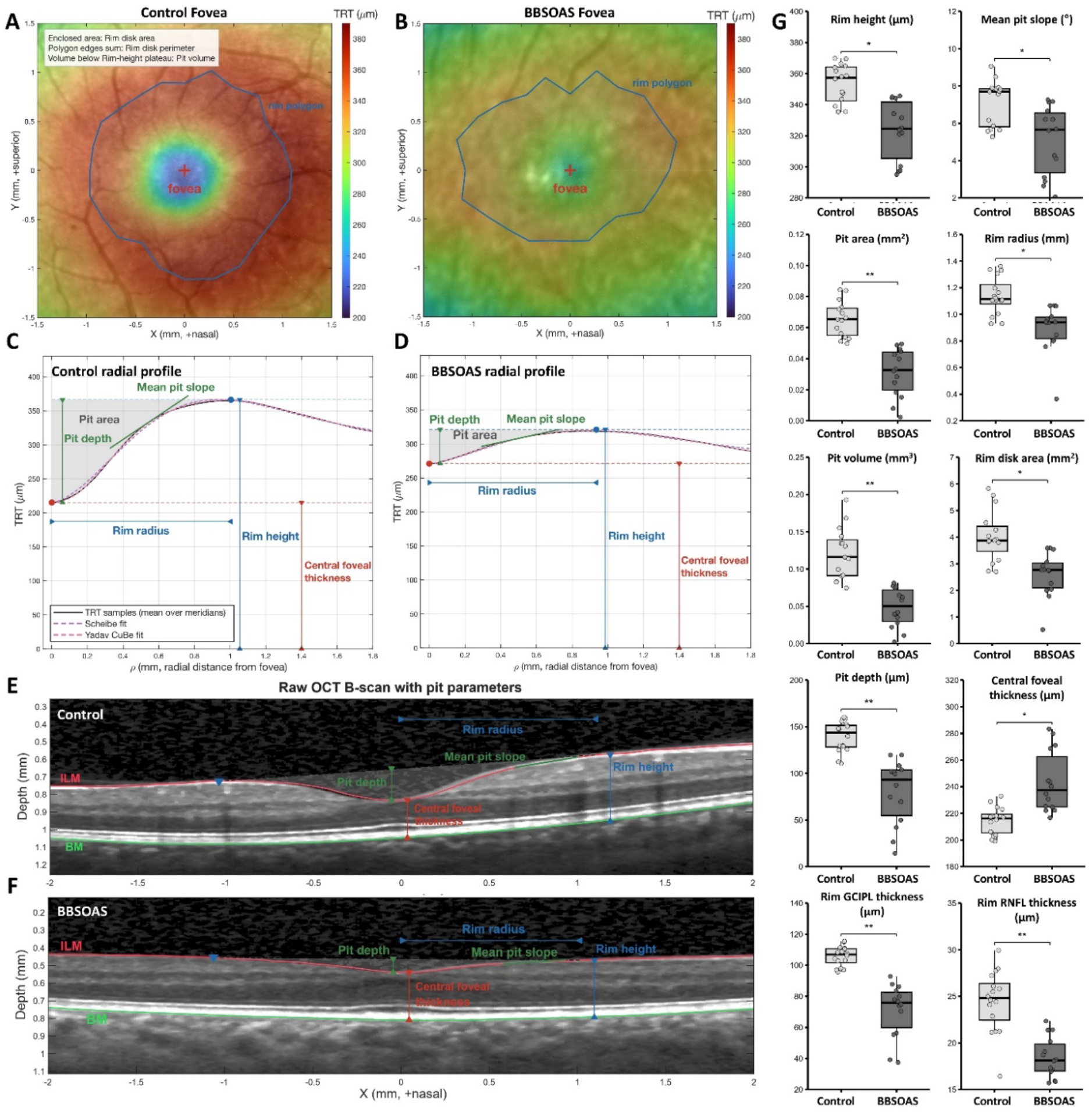
BBSOAS is associated with altered foveal morphology and hypoplasia. A,B) En-face total retinal thickness (TRT) maps overlaid on the corresponding infrared fundus images from representative eyes of a healthy control (A) and an individual with BBSOAS (B). A red cross indicates the fovea, and the blue polygon delineates the rim disk. **C,D)** Mean radial TRT profiles from the same representative control (left; C) and BBSOAS (right; D) eyes (black lines), with Scheibe (purple, solid) and Yadav cubic-Bezier (CuBe; magenta, dashed) parametric fits overlaid. Representative morphological parameters are indicated. **E,F)** Horizontal B-scans through the foveal center from the same representative control (E) and BBSOAS (F) eyes, with automated ILM (red) and BM (green) segmentation boundaries overlaid and representative morphological parameters annotated. The BBSOAS eye displays a shallower and broader foveal pit with increased central retinal thickness, consistent with foveal hypoplasia. **G)** Boxplots comparing selected foveal pit morphology parameters between individuals with BBSOAS (n = 14 eyes from 7 patients; dark grey) and age-matched healthy controls (n = 16 eyes from 8 individuals; light grey). Boxes represent the interquartile range, with the median indicated. Significance brackets show Benjamini–Hochberg-adjusted P values obtained from linear mixed-effects models (*p < 0.05, **p< 0.01, ***p < 0.001). The complete set of 22 quantified foveal morphology parameters is provided in *Fig. S9*. All values are expressed in millimeters for lateral dimensions and in micrometers for retinal thickness and pit depth. Demographic characteristics of patients with BBSOAS and control individuals included in the analysis are provided in *Table S1*, while *NR2F1* pathogenic variants in BBSOAS patients are listed in *Tables S2*. Summary statistics for the foveal parameters shown in the graphs are provided in *Table S3*.

To further characterize foveal morphology in three dimensions (3D), we applied complementary parametric modelling approaches to the OCT scans (*70*, *71*). Both models captured pronounced differences in pit shape features and highlighted striking differences between BBSOAS and control eyes (**Fig. 7C,D; Table S3**). The Scheibe asymptotic thickness α was reduced by more than half (0.03 ± 0.02 vs 0.08 ± 0.02 mm; *p* = 0.002), reflecting a thinner parafoveal retina. In the Yadav CuBe model, pit-bottom curvature β was also reduced (0.48 ± 0.12 vs 0.66 ± 0.10; *p* = 0.014), indicating a flatter pit and less sharply defined pit-rim transition, consistent with incomplete foveal excavation. At the foveal rim, inner retinal thickness was reduced in both the retinal nerve fiber layer (RNFL; 18.4 ± 2.2 vs 24.5 ± 3.3 µm; *p* = 0.002) and ganglion cell inner plexiform layer (GCIPL; 70.9 ± 17.4 vs 105.8 ± 6.4 µm; *p* = 0.002), consistent with parafoveal ganglion cell and axonal loss associated with optic neuropathy. Importantly, the central foveal phenotype is consistent with incomplete centrifugal migration of inner retinal layers, characteristic of foveal hypoplasia.

Interestingly, some of the affected parameters were consistent with a process causing retinal ganglion cell loss at the foveal rim, as expected in optic neuropathies, including autosomal dominant optic atrophy (ADOA) (**Tables S1, S4**) (*72*, *73*). These included changes in pit volume, depth and area, mean pit slope, and RNFL thickness. However, other foveal abnormalities, particularly increased central foveal thickness and foveal outer retinal thickness, were observed only in BBSOAS and are characteristic of foveal hypoplasia (complete data for all 22 foveal parameters are provided in **Fig. S9, Table S5**). Indeed, the reduction in rim height was less complete in BBSOAS than in ADOA, whereas pit size was similarly reduced, in part due to the increased foveal thickness. We therefore conclude that these changes cannot be fully explained by the generic consequences of optic nerve abnormalities, such as ganglion cell degeneration, associated with classical forms of optic neuropathy, and are more likely to reflect a BBSOAS-specific foveal pathological process.

Together, these clinical findings identify foveal hypoplasia as a previously unrecognized component of BBSOAS, providing evidence that *NR2F1* disruption affects not only the optic nerve and central visual pathways but also the developmental organization of the human central retina. This finding is consistent with our experimental data linking *Nr2f1*-dependent regulation of RA signaling and cone patterning to retinal specialization.

## DISCUSSION

High-acuity vision relies on the precise development of specialized retinal territories through the coordinated control of regional identity, progenitor competence, neuronal differentiation, and tissue morphogenesis (*66*). Although considerable progress has been made in defining the molecular mechanisms that control retinal neurogenesis, patterning and specialization (*2*, *66*, *74*), how disruption of these developmental programs contributes to human visual disorders remains poorly understood. By integrating single-cell transcriptomic analyses from three complementary *Nr2f1* mouse models with structural retinal imaging in individuals with BBSOAS, we identify *Nr2f1/NR2F1* as a key regulator of retinal, and particularly foveal specialization. We show that *Nr2f1* controls an RA-dependent transcriptional program by spatially regulating RA pathway genes, most notably *Cyp26a1*, thereby shaping retinal regional identity and cone photoreceptor distribution. Furthermore, we extend the evolutionary relevance of this regulatory mechanism to human NR2F1 biology by demonstrating direct binding of NR2F1 to conserved regulatory elements upstream of the *CYP26A1* locus. Importantly, the identification of reproducible foveal abnormalities in individuals with BBSOAS reveals a previously unrecognized component of NR2F1-associated visual pathology and provides a mechanistic framework linking disrupted retinal development to impaired visual function.

### NR2F1 regulates retinal patterning through a conserved RA-dependent developmental program

RA signaling provides a fundamental mechanism for retinal regionalization by generating spatially restricted transcriptional responses through tightly regulated RA synthesis and degradation (*66*, *75*). Although NR2F1 has been proposed to modulate RA-dependent transcription through interactions with RAR/RXR complexes and recruitment of transcriptional corepressors (*29*, *76*), whether this regulatory relationship operates during retinal development, and whether RA-related genes are directly regulated by NR2F1, has remained unclear. By comparing three independent *Nr2f1* mutant models, we identified a shared transcriptional signature enriched for RA pathway components, including *Crabp1, Crabp2, Rbp1, Cyp26a1,* and *Cyp26c1*. These findings demonstrate that distinct disease-associated *NR2F1* mutations converge on a common RA-dependent developmental program involved in retinal regionalization. Thus, RA signaling emerges as a central developmental axis downstream of *Nr2f1*, providing a potential mechanistic link between genetically diverse disease-causing mutations and retinal dysfunction, irrespective of mutation type. A major strength of this approach is the identification of a core *Nr2f1*-dependent network consistently dysregulated across distinct models, suggesting that perturbation of RA-related retinal patterning may represent a shared pathogenic mechanism across BBSOAS, rather than a mutation-specific effect.

Beyond providing *in vivo* evidence that *Nr2f1* regulates multiple RA pathway genes during retinogenesis, we identify *Cyp26a1* as a direct transcriptional target within this regulatory network. Loss of Nr2f1 function, through either gene deletion or patient-specific mutations, disrupted the spatial organization of *Cyp26a1* expression, resulting in an expansion of its dorso-equatorial domain into ventral retinal territories. *Cyp26a1* encodes an RA-degrading enzyme whose expression is itself induced by RA signaling, thereby contributing to a negative-feedback loop that locally limits RA availability (*54*, *77*, *78*). Interestingly, its homolog *Cyp26c1* has been reported to be largely unresponsive to exogenous RA administration in the mouse (*54*). Its dysregulation following *Nr2f1* loss or mutation therefore places Nr2f1 upstream of this regulatory network, rather than simply responding to changes in local RA levels. In addition, other components of the RA network were affected, including the ventral determinant *Vax2*, a regulator of *Cyp26a1* expression (*55*). Together, these findings indicate that *Nr2f1* contributes to the spatial restriction and maintenance of RA pathway activity during retinal development.

### Mutation-specific molecular consequences of *NR2F1* variants contribute to BBSOAS variability

The marked phenotypic variability observed among individuals with BBSOAS likely reflects, at least in part, differences in the molecular consequences of individual *NR2F1* variants (*34*). While gene deletions primarily reduce NR2F1 dosage (*31*), missense and truncating variants may additionally affect protein stability, subcellular localization, DNA binding, or interactions with transcriptional partners (*23*, *34*, *45*). Thus, although impaired NR2F1-dependent transcription represents a common consequence of NR2F1 dysfunction, mutation-specific effects are likely to further shape disease severity and clinical presentation (*34*).

Consistent with this view, the extent and nature of retinal molecular perturbations differed among the three *Nr2f1* mutant alleles. *R139L HET* retinae displayed the largest number of dysregulated genes, consistent with a strong impact of the DBD mutation on *Nr2f1*-dependent transcription (*32*, *33*, *36*). Increased *Ubc* expression, a component of the ubiquitin-proteasome system, in both *R139L HET* and *HOM* retinae (**Fig. S4**), together with reduced Nr2f1 protein despite increased transcript levels, is consistent with altered protein stability and/or enhanced protein turnover. Interestingly, the truncated E397* protein, abnormally localized in the cytoplasm, was even more strongly reduced than the R139L protein, indicating that this LBD-truncating mutation affects both protein abundance and subcellular distribution. Whether these differences reflect distinct protein quality-control mechanisms or contribute to dominant-negative or gain-of-function effects remains to be determined.

Mutation-specific effects were also evident at the transcriptional level. Gene ontology analysis further revealed distinct mutation-specific differences, with *Null*, *E397\**, and *R139L* mutants preferentially affecting processes related to apoptosis/neurogenesis, broader developmental programs, and cell migration/axon guidance, respectively (**Fig. S5**). Similarly, although all three models shared a core RA-related transcriptional signature, additional pathway components were selectively dysregulated in individual alleles. For example, *Aldh1a3*, *Aldh1a7*, as well as the RA receptor gene RARα, were affected in *E397\** and *R139L* retinae, whereas the magnitude of dysregulation of shared genes also varied between alleles, with *Crabp1* most strongly affected in *R139L* and *Cyp26c1* particularly upregulated in *E397\** mutants. Thus, the DBD mutation did not uniformly produce the strongest downstream effects, highlighting the complexity of allele-specific transcriptional consequences.

Together, these findings demonstrate that *NR2F1* variants are not functionally equivalent. Rather, their effects reflect a combination of altered protein stability, subcellular localization, DNA-binding capacity, and downstream transcriptional regulation. This molecular heterogeneity is likely to contribute to the genotype-phenotype relationships underlying the remarkable clinical variability of BBSOAS and emphasizes the importance of considering the specific molecular consequences of individual *NR2F1* variants when interpreting disease mechanisms.

### *NR2F1*-dependent RA signaling links retinal patterning to foveal specialization and visual function

RA gradients are important regulators of photoreceptor subtype specification (*59*). Consistent with this role, we found that *Nr2f1*-dependent regulation of RA signaling influences cone photoreceptor patterning, a key determinant of retinal function. Loss of *Nr2f1* or expression of patient-specific mutations altered the spatial distribution of S-and M-opsin-expressing cones along the D-V axis, further implicating Nr2f1 in cone subtype specialization (*79*). Such alterations in cone specification or distribution could contribute to the reduced visual acuity and color vision abnormalities reported in some individuals with BBSOAS (*26*, *34*, *38*).

In humans and other diurnal species, cone photoreceptors reach their highest density and acquire a highly specialized spatial organization within the fovea, the region of the primate retina responsible for high-acuity and color vision (*4*). Although mice lack a morphologically defined fovea, studies in chick and primates indicate that conserved RA-dependent mechanisms contribute to the formation of cone-enriched retinal territories (*20*, *54*, *56*, *69*, *74*). Across species, the RA-degrading enzyme CYP26A1 is enriched in developing high-acuity retinal regions (*69*, *74*), suggesting that spatial restriction of RA signaling represents a conserved developmental strategy for retinal specialization (*66*). More recently, CYP26A1 has been shown to play an essential role in cone specification and foveal identity in human fetal tissue and retinal organoids (*56*, *67*), and *CYP26A1* variants have been identified as a cause of foveal hypoplasia (*80*). Despite this evolutionarily conserved role, the upstream mechanisms that establish the highly restricted expression of *CYP26A1* and the associated RA-depleted foveal domain have remained poorly understood (*66*). Our findings identify Nr2f1/NR2F1 as an upstream regulator of this conserved developmental program, controlling the spatial distribution of RA activity through regulation of *Cyp26a1* and other RA pathway genes. By shaping the spatial domains and cellular levels of RA signaling during embryonic retinogenesis, Nr2f1/NR2F1 may establish the transcriptional environment required for retinal regional specialization, including appropriate photoreceptor subtype specification and, in species possessing one, foveal development. In line with the altered foveal morphology identified in BBSOAS patients, we propose NR2F1 as a novel foveal hypoplasia-associated gene.

This model provides a potential mechanistic explanation for the foveal abnormalities identified in BBSOAS. Although the mouse retina does not contain a fovea, several genes showing D-V regionalization in our study are also differentially expressed along the central (foveal)-to-peripheral retinal axis in diurnal primates (*81*). Notably, genes enriched in the ventral mouse retina, such *Vax2* and *Nr2f1* itself, are preferentially expressed in the peripheral primate retina, whereas dorsal genes such as *Cyp26a1* are enriched in the foveal region (*81*). This correspondence raises the possibility that developmental programs originally involved in D-V patterning in mice have been evolutionarily co-opted to regulate central-peripheral specialization in species with cone-enriched, high-acuity retinae. In this framework, the complementary expression of *NR2F1* and *CYP26A1* identified in the mouse retina may reflect a conserved regulatory architecture that contributes to the formation of specialized retinal territories across species. The reproducible structural foveal abnormalities in BBSOAS provides the first clinical evidence supporting this model. The combination of altered foveal morphology in BBSOAS, conserved expression of *CYP26A1* in the developing primate fovea, and the role of NR2F1 in regulating *Cyp26a1* in the mouse retina suggests that disruption of an *NR2F1*-dependent RA regulatory program may contribute to abnormal foveal development and visual dysfunction. Importantly, our data do not yet establish a direct regulation of *CYP26A1* expression by NR2F1 during human foveal development. This will require additional experimental models, such as human retinal organoids and other stem cell-derived models that recapitulate key aspects of foveal development. Nevertheless, our findings provide a mechanistic framework linking *NR2F1* dysfunction to impaired retinal specialization and suggest that modulation of CYP26A1 activity, or more broadly of RA signaling, could represent a potential therapeutic strategy. Whether such approaches can correct developmental retinal defects will require experimental validation.

The identification of foveal abnormalities also broadens the understanding of visual dysfunction in BBSOAS. Although the syndrome was initially characterized as an optic atrophy disorder (*31*), visual impairment is increasingly recognized as a complex phenotype involving multiple components of the visual system, including optic nerve abnormalities and cortical visual impairment (*25*, *26*, *34*, *39*). Reduced visual acuity is among the most consistent clinical features (*32*), yet its developmental basis has remained incompletely understood. Our findings provide evidence for a retina-intrinsic component of BBSOAS visual dysfunction, extending the disease spectrum beyond the optic nerve and central visual pathways. In particular, the presence of foveal abnormalities suggests that impaired retinal specialization may contribute to reduced visual acuity in BBSOAS and highlights retinal development as an important, previously underappreciated component of *NR2F1*-associated visual pathology.

### Limitations and perspectives

Several considerations should be taken into account when interpreting our findings. The three *Nr2f1* mouse models examined here represent only a subset of the genetic variation observed in BBSOAS and therefore cannot be generalized to all DBD, LBD, or other *NR2F1* variants. Nevertheless, the transcriptional changes shared across all three models identify a core signature that may represent a common mechanism underlying different mutation types. In addition, our scRNA-Seq analyses were performed in mouse, whose retina lacks a fovea, limiting the extent to which the identified mechanisms can be directly extrapolated to human foveal development. Human retinal organoids, *ex vivo* fetal retinal tissue, and other models with fovea-like specializations will be important to determine whether the NR2F1-CYP26A1 axis is conserved in human retinal specialization. Finally, our analysis focused on retinal progenitor cells at a single developmental stage and therefore provides only a snapshot of NR2F1-dependent molecular changes. Analyses of additional retinal cell types and developmental stages will be needed to define the broader cell type-and stage-specific functions of *NR2F1*. Despite these limitations, the convergence of distinct *NR2F1* mutations on a shared RA-related transcriptional program, together with mutation-specific differences in its regulation, provides a framework for understanding both common and variable features of BBSOAS-associated visual dysfunction.

## MATERIALS AND METHODS

### Ethical statements

All mouse experiments and standard housing conditions were in accordance with relevant national and international guidelines and regulations (European Union Directive 2010/63/EU), under authorization from the French Ministry for Higher Education and Research, and with approval from the local ethics committee in France (CIEPAL Azur 28; NCE/2024-976). The work on human patients was approved by the research study “Genotype and Phenotype in Inherited Neurodegenerative Diseases (REC reference: 13/YH/0310)” from the Cambridge University Hospitals NHS Foundation Trust.

### Animal housing

Briefly, adult mice were housed under a 12-h light/dark cycle, with up to three animals per cage, provided with standard environmental enrichment (wooden cubes, cotton pads, and igloos), and given *ad libitum* access to food and water. *Nr2f1 knock-out* (*KO*) or *knock-in* (*KI*) mice were generated and genotyped to obtain heterozygous (*HET*) and homozygous (*HOM*) embryos and animals, as previously described (*28*, *82*). Littermates of *HET* and *HOM* mice carrying normal *Nr2f1* alleles were used as *wild-type* controls (*WT*). Midday of the day of vaginal plug detection was designated as embryonic day 0.5 (E0.5). Control and mutant mice were maintained on a C57BL/6J background. Both male and female embryos and post-natal mice were used in this study; the age of each specimen is indicated in the corresponding experiments.

### Real time qRT-PCR

Mouse embryo brains and eyes were dissected on ice-cold PBS and immediately frozen at-80 °C. Total RNA was isolated using NucleoSpin RNA II columns (Macherey-Nagel, 740902.50). RNA quantity and quality were assessed using a NanoDrop spectrophotometer (Thermo Fisher Scientific) and agarose gel electrophoresis. For each sample, 200 ng of total RNA was reverse-transcribed using random nonamers (Reverse Transcriptase Core Kit, Eurogentec, qRTRTCK-03). Quantitative reverse transcription PCR (qRT-PCR) was performed using GoTaq SYBR Green qPCR Mix (Promega, A6001) on a LightCycler® 480 system (Roche). cDNA samples were stored at-20 °C and diluted to obtain 2 ng of cDNA per reaction. Amplification take-off values were determined using the built-in LightCycler® 480 relative quantification analysis software, and relative gene expression levels were calculated using the 2^-ΔΔCt method, with normalization to the housekeeping gene *Actin*. At least two technical replicates were performed for each sample and gene analyzed. Primer sequences used were as follows: *Nr2f1* exon 3, forward 5′-TCCCATCGAAACTCTCATCC-3′ and reverse 5′-AGTGGGCTGCTCTTGTTCC-3′; *Nr2f1* R139L WT, forward 5′-CAACCAGTGCCAATACTGCCGCC-3′ and reverse 5′-CAGACAGGTAGCAGTGGCCA-3′; *Nr2f1* R139L KI, forward 5′-CAACCAGTGTCAGTATTGTCT-3′ and reverse 5′-TAGCCAGACAGGTAGCAGTG-3′; *Nr2f1* E397* KI, forward 5′-AAGCCAGTACCCCAACCAG-3′ and reverse 5′-AACATATCTCGGATGAGAGTTCAT-3′; *Nr2f1* universal control primers, forward 5’-CGAGGGCTGCAAAAGTTTCT-3’ and reverse 5’-GCATTCTTCCTCGCTGAACC-3’; *Actin*, forward 5′-ATGTGGATCAGCAAGCAGGA-3′ and reverse 5′-GTGTAAAACGCAGCTCAGTAACA-3′.

### Immunostaining

Basic procedures for brain tissue fixation and cryostat sectioning are described in detail in a previous methodological report (*83*). Retinae were collected in 2-ml Eppendorf tubes and fixed in 2 ml of 4% paraformaldehyde (PFA) at 4 °C for 3-4 h under gentle agitation. Samples were then washed twice in 1× PBS and cryoprotected by incubation in 10% sucrose (Sigma-Aldrich, S9378-1KG; dissolved in 1× PBS) for a minimum of 4 h up to overnight, followed by 25% sucrose (dissolved in 1× PBS) overnight at 4 °C. After cryoprotection, most of the sucrose solution was removed and replaced with OCT compound (Leica Tissue Freezing Medium, 14020108926, or Cryomatrix, 6769006) for 10 min under gentle rotation. Samples were then transferred to embedding molds filled with fresh OCT and stored at-80 °C. Cryosections (12 μm thick) were cut using a Leica CM3050S cryostat and collected onto Superfrost Plus glass slides (Thermo Fisher Scientific, J1800AMNZ, or VWR, 631–0108). Slides were stored at-80 °C until use. Prior to immunostaining, sections were air-dried at room temperature for 10-15 min and washed twice in 1× PBS (10 min each) to remove residual OCT. Heat-induced antigen retrieval was performed in 0.1 M sodium citrate buffer (pH 6.0) at 95 °C for 10 min. After a 5-min wash in 1× PBS to remove the unmasking solution, sections were incubated in blocking solution for 1 h at room temperature. The blocking solution consisted of 1× PBS containing 5% serum (sheep serum, Sigma, S2263-100ML; or newborn calf serum, Thermo Fisher Scientific, 16010-167, for primary antibodies raised in sheep or goat) and 0.3% Triton X-100 (Sigma-Aldrich, T8787-250ML). Primary antibodies were incubated for a minimum of 4 h at room temperature or overnight at 4 °C in 1× PBS supplemented with 1% serum and 0.1% Triton X-100. Primary antibodies used in this study were: rabbit anti-NR2F1 (Abcam, ab181137; 1:1000), sheep anti-CHX10/VSX2 (Abcam, ab16141; 1:500), rabbit anti-VAX2 (Proteintech, 15773-1-AP; 1:500), rabbit anti-CRABP2 (Proteintech, 10225-1-AP; 1:500), rabbit anti-βIII-tubulin (TUBB3) (BioLegend, 802001; 1:1000), rabbit anti-S-opsin (Sigma, ABN1660-I; 1:1000), and rabbit anti-M-opsin (red/green opsin) (Millipore, AB5405; 1:1000). Alexa Fluor 488, 555, 594, and 647-conjugated secondary antibodies (anti-mouse, anti-rabbit, anti-rat, anti-goat, anti-guinea pig, or anti-sheep IgG; Thermo Fisher Scientific) were used at a dilution of 1:500 and incubated for 2 h at room temperature or overnight at 4 °C. The secondary antibody solution also contained DAPI (1:1000; Thermo Fisher Scientific, 10374168) for nuclear staining. Following final washes (3x times for 10 min each in 1× PBS), sections were mounted with mounting medium (80% glycerol, 2% n-propyl gallate in 1× PBS), coverslipped, and sealed with nail polish. Stained sections were stored at-20 °C. Images were acquired using a Zeiss Apotome microscope with AxioVision software and exported as TIFF files. Alternatively, images were collected using a Vectra Polaris slide scanner (Akoya Biosciences) and analyzed using AI-assisted image analysis with HALO software (Indica Labs).

### *In situ* hybridization

Embryos used for in situ hybridization (ISH) were fixed overnight in 4% paraformaldehyde (PFA) at 4 °C, cryoprotected in 25% sucrose, embedded in OCT compound, stored at-80 °C, and sectioned at 14 μm using a Leica cryostat. Digoxigenin (DIG)-labeled RNA probes for Sfrp2 and Crabp1 were generated from mouse retinal cDNA using gene-specific primers containing a T7 RNA polymerase promoter sequence (see below). The Vax2 probe was generated by plasmid linearization with HindIII followed by in vitro transcription with T3 RNA polymerase from a plasmid kindly provided by P. Bovolenta (*49*). Amplified cDNA fragments were purified, and concentration and purity were assessed by spectrophotometry. *In vitro* transcription was performed using the DIG RNA Labeling Mix (Roche) according to the manufacturer’s instructions. RNA probes were purified by LiCl/ethanol precipitation, resuspended in nuclease-free water, verified by agarose gel electrophoresis, and stored at-20 °C until use. Head cryosections were thawed, air-dried for at least 1 h at room temperature, washed twice in 1× PBS, and permeabilized by two consecutive 10-min incubations in RIPA buffer (150 mM NaCl, 1% NP-40, 0.5% sodium deoxycholate, 0.1% SDS, 1 mM EDTA, 50 mM Tris-HCl, pH 8.0). Sections were post-fixed in 4% PFA for 15 min, washed three times in PBS, incubated in 3% hydrogen peroxide for 30 min at room temperature, acetylated for 15 min in freshly prepared 100 mM triethanolamine containing 0.25% acetic acid, and washed three times in PBS. Pre-hybridization was performed for 1 h at room temperature using pre-warmed hybridization buffer containing 50% formamide, 5× saline sodium citrate (SSC), 5× Denhardt’s solution (Invitrogen), 500 μg/mL salmon sperm DNA (Ambion), and 250 μg/mL yeast tRNA. DIG-labeled RNA probes were denatured at 90 °C for 5 min and added to the hybridization solution at a final concentration of 1 μg/mL. Hybridization was carried out overnight at 65 °C (or 68 °C for selected probes) in a humidified chamber containing 50% formamide and 5× SSC. Following hybridization, sections were subjected to stringent washes at 65

°C in 50% formamide, 5× SSC (pH 4.5), 1% SDS, followed by 50% formamide, 2× SSC (pH 4.5), and 0.2% SDS. Sections were then equilibrated in MABT buffer (100 mM maleic acid, 150 mM NaCl, 0.1% Tween-20, supplemented with 0.5 mg/mL tetramisole hydrochloride), blocked sequentially in MABT containing Roche blocking reagent and subsequently in MABT supplemented with 10% sheep serum. Hybridized probes were detected by overnight incubation at 4 °C with alkaline phosphatase-conjugated anti-digoxigenin antibody (Roche/Sigma, 1:1,000) diluted in MABT containing 10% sheep serum. Sections were washed extensively in MABT, equilibrated in B3 buffer (100 mM Tris-HCl, pH 9.5, 50 mM MgCl₂, 100 mM NaCl, 0.1% Tween-20, 0.5 mg/mL tetramisole hydrochloride), and developed using NBT/BCIP or BM Purple substrate (Roche) supplemented with 0.1% Tween-20 at room temperature or 37 °C until the desired signal intensity was reached. Color development was stopped by washing in PBS containing 0.1% Tween-20. Sections were briefly rinsed in distilled water, mounted with Aquatex mounting medium (Sigma), dried overnight at room temperature, and stored until imaging. The primers used to amplify target sequences and produce RNA probes used in this study were: *Sfrp2*, probe size: 579 bp, Forward primer: 5’-CCAAGAAGTTCCTGTGCTCG-3’, Reverse primer with T7 sequence in lowercase: 5’-taatacgactcactatagggcTGAACTCTCTCTCTGGCCCTTC-3’; *Crabp1*, probe size: 559 bp, Forward primer: 5’-GCCACCATGCCCAACTTC-3’, Reverse primer with T7 sequence in lowercase: 5’-taatacgactcactatagggcTGTCATGGGGAAAATGGGGT-3’.

### RNAscope hybridization

RNAscope *in situ* hybridization was performed on cryosections of E14.5 mouse embryonic eyes using the RNAscope Multiplex Fluorescent Reagent Kit v2 (Advanced Cell Diagnostics, ACD, Newark, CA, USA) according to the manufacturer’s instructions. Briefly, embryonic heads were fixed, cryoprotected, embedded in OCT, cryosectioned, and mounted onto Superfrost Plus microscope slides. Tissue sections were pretreated following the standard RNAscope protocol and hybridized with probes specific for mouse Cyp26a1 (Catalog # 468911) and Cyp26c1 (Catalog # 481861) (ACD). Following signal amplification, probe detection was performed using ClariTSA™ Fluorophore Kit 570 (Catalog # 7526) and ClariTSA™ Fluorophore Kit 650 (Catalog # 7527) (Tocris-Bio-Techne). Nuclei were counterstained with DAPI and slides were mounted with Fluoromount-G™ (00-4958-02; Invitrogen-Thermo Fisher Scientific). For combined RNAscope and IF, RNAscope™ Manual PretreatPro™ (Catalog # 323770; ACD) was used in place of RNAscope™ Protease Plus (Catalog # 322331; ACD) for tissue permeabilization to preserve antigenicity. Probe signals were detected using the Opal™ 520 Reagent Pack (FP1487001KT; PerkinElmer) and Opal™ 570 Reagent Pack (FP1488001KT; PerkinElmer), or Opal™ 650 Reagent Pack (FP1496001KT; Akoya Biosciences), depending on the experimental design. Fluorescence images were acquired using either a Zeiss Apotome microscope or a Vectra Polaris Automated Quantitative Pathology Imaging System (Akoya Biosciences). Within each experiment, all samples were imaged using identical microscope settings, including laser power or illumination intensity, exposure time, detector gain, and optical configuration, to enable direct comparison of fluorescence signal intensities.

### Statistical analysis

All data were statistically analyzed and graphically represented using Microsoft Office Excel or GraphPad Prism (versions 7.00 and 8.00). Quantitative data are presented as mean ± SEM. For cell percentage or cell number quantification following immunostaining, measurements were performed on at least three sections derived from two to three different embryos, unless otherwise stated. Sections displaying damaged histology were excluded from further analysis. Microscopy images were processed using Photoshop or ImageJ software for cell counting, as indicated in the figures and figure legends. When calculating percentages relative to total cell number, totals were determined by counting DAPI-positive nuclei, unless otherwise specified. Statistical analyses were performed using one-way or two-way ANOVA, depending on the number of independent variables, with Tukey’s correction. Statistical significance was defined as follows: * p < 0.05; ** p < 0.01; *** p < 0.001; **** p < 0.0001.

### Extract preparation for native holdup (nHU)

For validation experiments, we used day 12 (d12)-differentiated neuronal progenitor stem cells (NPSCs), and extracts were prepared using the same protocol as described below. For fluorescence-based measurements, the pEGFP-C1-NR2F1 (canonical sequence, UniProt #P10589) mammalian expression plasmid was generated by standard restriction enzyme cloning. Point mutations were introduced using the standard QuikChange site-directed mutagenesis protocol with Platinum SuperFi II DNA Polymerase (Thermo Fisher Scientific). HEK293T cells (authenticated by STR profiling as a 100% match to the ATCC reference CRL-3216, RRID: CVCL_0063, and confirmed to be mycoplasma-free prior to the experiments using the Eurofins Mycoplasma test) were cultured in Dulbecco’s Modified Eagle Medium (DMEM; Gibco, 1 g/L glucose) supplemented with 10% fetal calf serum (FCS) and 1% Penicillin-Streptomycin (Gibco) in CellBIND flasks (Corning). Cells were passaged at a 1:10 ratio every 3-4 days. For transient transfection, cells were seeded onto poly-D-lysine-coated (Sigma-Aldrich, P6407) 100 mm culture dishes at a density of 2×10^6^ cells/dish. The following day, HEK293T cells were transfected with the pEGFP-C1-NR2F1 plasmid using JetPRIME reagent according to the manufacturer’s instructions. Cell extracts were prepared as previously described (*62*, *63*).

### Native holdup (nHU) titrations to measure binding of NR2F1 to the *CYP26A1* regulatory region

Native holdup (nHU) titration experiments were performed as previously described (*62*). To validate the activity of the GFP-tagged NR2F1 protein relative to the native wild-type protein extracted from human neural progenitors, we used the human version of the previously described *RND2* recognition sequence (*64*). The nHU assay was then used to assess NR2F1 binding to the human *CYP26A1* regulatory region. A random DNA sequence was included as a negative control. All DNA oligonucleotides were synthesized by Eurofins Genomics (Eurofins Genomics Germany GmbH, Germany) and purified by high-performance liquid chromatography (HPLC). The following oligonucleotide sequences were used (predicted NR2F1 recognition sites are shown in **bold**): **RND2_BIOT_F**, BioTEG-acctc**TGACC**TC**TGACCT**gatcc; **RND2_REV**, ggatc**AGGTCA**GA**GGTCA**gaggt; **CYP26A1_BIOT_F**, BioTEG-TGCAGGGGTCACAGGCG**GGTCAG**GCCCGAGGATTGGGAATGGGCCCC**AGGTCAC**GT CC; **CYP26A1_R**, GGACG**TGACCT**GGGGCCCATTCCCAATCCTCGGGCC**TGACCC**GCCTG**TGACCC**CTG CA; **RANDOM_BIOT_F**, Biotin-CCACTCTGTTCTAGAGAAGCCTTCGTGAACT; **RANDOM_R**, AGTTCACGAAGGCTTCTCTAGAACAGAGTGG.

The assay was performed in 96-well filter plates with a starting DNA bait concentration of 5 uM in a 12-point titration series. After incubation of the oligo-saturated resins with cellular extracts, the supernatant was quickly separated and the unbound fraction was determined from these supernatant solutions. Using the equilibrium binding equation, the bound fractions could be converted to apparent binding affinities. Fluorescence measurements were performed directly on the samples using a SpectraMax ID5 plate reader (Molecular Devices). Each data point was measured in triplicate. The data were fitted using the Hill equation (cooperative binding model) in QtiPlot, as this model provided the best fit for multiple datasets obtained using NR2F1 extracts.

To assess NR2F1 quantity in endogenous nHU measurements, a Trans-Blot Turbo Transfer System and a Trans-Blot Turbo RTA Midi 0.45 μm LF PVDF Transfer Kit (ref. 1704275; Bio-Rad) were used. The following primary antibodies were used: anti-NR2F1 antibody (1:5,000, overnight 4 °C, Abcam ab181137) and anti-glyceraldehyde-3-phosphate dehydrogenase (GAPDH) primary antibody (1:5,000, 1 hr RT, ref. MAB374; Sigma-Aldrich, RRID:AB_2107445). The following secondary antibodies were used: Jackson ImmunoResearch, West Grove, PA, USA; peroxidase-conjugated AffiniPure goat anti-rabbit (H + L), ref. 111-035-003 and anti-mouse, ref. 115-035-146 (RRID:AB_2313567 and RRID:AB_2307392). For detection, a chemiluminescent horseradish peroxidase substrate (ref. WBKLS0100; Immobilon, MilliporeSigma, Burlington, MA, USA) was used and blots were imaged using an Amersham ImageQuant 600 imaging system.

### Single cell RNA sequencing

#### Cell dissociation for single-cell RNA sequencing and flow cytometry analysis

E14.5 embryos were collected in ice-cold 1× PBS, and retinae with surrounding tissues were dissected and placed in low-bind Eppendorf tubes to minimize material loss. A small tissue sample from each embryo was collected separately for genotyping. Retinae were dissociated in Trypsin at 37 °C for a total of 25 min. During dissociation, samples were gently mixed every 5 min: the first two rounds consisted of tube inversion, followed by gentle pipetting in subsequent rounds to promote single-cell dissociation. Dissociated cells were filtered through a 40 μm nylon cell strainer (FALCON, 352340), centrifuged at 1200 rpm (≈170 × g) for 5 min, and resuspended in 1× PBS supplemented with 0.5% BSA (Jackson, 001-000-161). Cell viability was assessed by Trypan Blue staining prior to counting for library preparation. Samples were considered suitable for single-cell RNA sequencing when ≤5% of cells were damaged. Cells were fixed using the Library Prep RNA Fixed_4n Single Cell protocol (10X Genomics) according to the manufacturer’s instructions and stored at-80 °C for up to 4 months before processing.

#### Single-cell RNA sequencing: library preparation and sequencing

Single-cell RNA sequencing was performed using 10X Genomics technology. Library preparation was conducted at the GenomEast platform (Institute of Genetics and Molecular and Cellular Biology) using the Chromium Fixed RNA Profiling Reagent Kits for Multiplexed Samples (PN CG000527). Single-cell libraries were generated with the Chromium Next GEM Single Cell Fixed RNA reagent set (10X Genomics, Leiden, The Netherlands) following the manufacturer’s instructions. Cell counting was performed using an automated Luna FX7 counter (Logos Biosystems, South Korea). For each of the 12 samples (*WT, HET, HOM* embryos from each mouse line with double sequencing of *E397\** samples), 200000-300000 starting cells were hybridized with a unique Mouse WTA Probe Barcode (10X Genomics, PN-1000497) in 80 µL of hybridization mixture plus 20 µL of WTA probes at 42 °C for 19.5 h. After hybridization, equal numbers of cells from each reaction were pooled, washed, and recounted. Cells were then loaded onto the Chromium X using a Next GEM Chip Q, targeting 5000 cells per hybridization reaction, for a total of 120000 cells per 12-plex library. Full-length cDNA amplification was performed using 8 PCR cycles, followed by dual-index library construction with 8 cycles during sample index PCR. Library quality and quantity were assessed using a Bioanalyzer 2100 (Agilent Technologies, Les Ulis, France). Libraries were sequenced on an Illumina NextSeq 2000 as paired-end 28 + 85 bp reads. Image analysis and base calling were performed using RTA version 2.7.7 and BCL Convert version 3.8.4.

#### Analysis of differentially expressed genes (DEGs) and gene enrichment

Raw counts were loaded into the Seurat R package (version 4.4.0) (*84*). Counts were normalized using the NormalizeData function with default parameters (LogNormalize method). Differential gene expression analysis between *HOM* and *WT* (*WT versus HOM* DEGs) or between *HET* and *WT* (*WT versus HET* DEGs) within each sample was performed using the FindMarkers function on normalized counts. Only genes detected in at least 5% of cells (minimum of three cells) in either population were tested. P values were calculated using the Wilcoxon rank-sum test and adjusted using the Bonferroni correction.

#### Gene ontology and over-representation analyses

Differentially expressed genes (DEGs) identified by Seurat were further processed using a custom Python (version 3.11.9) analysis pipeline. Prior to downstream analyses, genes annotated as deprecated Ensembl entries and Y chromosome genes were removed to avoid annotation inconsistencies and potential sex-related bias. Gene annotations were retrieved from the Ensembl BioMart database (GRCm39, Mus musculus). DEGs were defined using an adjusted P value < 0.05 together with an absolute average log2 fold change (|avg_log2FC|) > 0.1. This threshold was selected to match the reference analysis while retaining biologically relevant transcriptional changes, including *Nr2f1*. Gene Ontology (GO) over-representation analysis (ORA) was performed using the Python package gseapy (version 1.2.1), which queries the Enrichr database. Enrichment analyses were carried out independently for each genotype using the complete DEG lists, as well as genotype-specific and shared DEG subsets identified by Venn diagram analysis. Gene sets were interrogated against the GO Biological Process 2023, GO Molecular Function 2023, GO Cellular Component 2023, and KEGG 2019 Mouse databases. Disease enrichment analyses were additionally performed using the DisGeNET database. Enriched terms with an adjusted P value < 0.05 were considered statistically significant. To identify molecular signatures associated with specific *Nr2f1* alterations, DEGs were partitioned into genotype-exclusive and shared gene sets based on their overlap among the *Null, E397*,* and *R139L* models. ORA was performed separately for each Venn diagram subset, including the intersection common to all three genotypes, allowing the identification of mutation-specific and shared biological pathways. Gene sets containing fewer than three genes were excluded from enrichment analyses. In addition to enrichment analyses, overlap and concordance between DEG datasets were assessed using Venn diagrams (matplotlib-venn, version 1.1.2), UpSet plots (upsetplot, version 0.9.0), pairwise log2 fold-change correlations (scikit-learn, version 1.8.0), principal component analysis (PCA), and heatmaps or dot plots of shared DEGs generated with the custom Python pipeline.

#### Dot plot visualization of gene expression

Dot plots were generated using a custom R script (Seurat, ggplot2, and patchwork packages) to visualize the expression of selected genes across genotypes. For each gene, the percentage of expressing cells and the average normalized expression were extracted from the Seurat object for each sample (*WT*, *HET*, and *HOM*) within each of the three *Nr2f1* lines. Dot size represents the percentage of cells expressing the gene, whereas color intensity reflects the log2 fold change in average normalized expression relative to the *WT* reference of the same line. Figures were exported in both PDF and PNG formats using the ggplot2 package.

#### Pseudo-spatial annotation of retinal progenitor cells along the dorso-ventral and naso-temporal axes

To investigate regional transcriptional heterogeneity within the retinal progenitor cell (RPC) population, RPCs were assigned continuous pseudo-spatial coordinates along the dorso-ventral (D-V) and naso-temporal (N-T) retinal axes using a custom Seurat pipeline. Raw counts were log-normalized using the Seurat Normalize Data function. Regional identities were assigned based on the expression of five categories of established topographical marker (anchor) genes: nasal (*Foxg1*), temporal (*Foxd1*), ventral (*Nr2f1*, *Aldh1a3*, *Vax1*, and *Vax2*), dorsal (*Bambi*, *Aldh1a1*, *Tbx5*, *Tbx3*, *Tbx2*, and *Nr2f2*), and equato-dorsal (*Cyp26a1* and *Cyp26c1*). These positional marker genes enabled the identification of spatially distinct RPC subpopulations, which were subsequently used for downstream differential expression and pathway analyses. For each anchor gene, log-normalized expression was calibrated against its 99th percentile expression level in the wild-type reference sample (*Null_WT*, sample SWRN1), and calibrated values were averaged within each category to obtain per-cell dorsal, ventral, nasal, temporal, and equato-dorsal scores. D-V and N-T scores were then computed as (Dorsal - Ventral) and (Nasal - Temporal), respectively. A two-dimensional reference space was built by principal component analysis (PCA) using the four canonical-axis anchor gene categories (Dorsal, Ventral, Nasal, and Temporal), restricted to wild-type reference cells, and oriented to align with the D-V and N-T marker scores. All cells were then projected onto this fixed reference geometry and rescaled into a bounded two-dimensional coordinate system (x-axis, temporal-to-nasal; y-axis, ventral-to-dorsal) based on each cell’s distance rank within the wild-type reference distribution. Because *Cyp26a1* and *Cyp26c1* mark a band located dorsal to the retinal equator by *in situ* hybridization (*20*, *54*), cells with higher expression of these markers had their D-V coordinate adjusted toward this expected position, proportionally to their expression level, without altering their N-T coordinate. For each of the nine biological groups, the mean pseudo-spatial position of all cells was plotted, and a vector was drawn from the wild-type reference centroid to each of the other group centroids to visualize genotype-associated positional shifts. The spatial expression patterns of *Cyp26a1* and *Cyp26c1* were visualized by mapping each cell log-normalized expression onto its pseudo-spatial coordinates, separately for each biological group.

#### Foveal pit morphology analysis

Macular spectral-domain OCT volumes (Heidelberg Spectralis) were exported and processed using a custom modified version of the RETIMAT toolbox1. Retinal layer thicknesses were derived from the automated segmentation boundaries. The foveal center was localized on the retinal map using a flood-fill algorithm constrained within a 1.5 mm radius of the scan center. Segmentations were resampled onto a radial grid with 24 meridians, and pit morphology was extracted per-meridian: central foveal thickness (CFT), pit depth, pit area (the cross-sectional area between the rim plateau and the retinal curve), rim height, rim radius, mean pit slope, rim disk area/perimeter, and pit volume. Two recently described parametric models were then fitted to the radial TRT profile (*70*, *71*): the Scheibe model yielding rate (μ), rim-prominence (σ), shape (γ) and asymptotic-thickness (α) coefficients, and the Yadav CuBe model, yielding interior and exterior Bézier control-point tensions (α_int, β, α_ext) and shoulder coordinates (P2x, P2y). Retinal nerve fiber layer (RNFL) and ganglion cell layer (GCL) thicknesses were sampled at each meridian’s rim position; foveal inner and outer retinal thicknesses were measured at the foveal center. All per-meridian values were averaged across all meridians. Statistical testing was performed in R. Both eyes were included in analyses, with within-patient correlation modelled by a random intercept in a linear mixed-effects model. Benjamini-Hochberg adjustment was used to correct for multiple comparisons.

## Acknowledgments

The authors thank the Experimental Histopathology facility, the PRISM microscopy facility, particularly Sameh Ben Aicha and Mathis Courtillet, and the animal facility, particularly Kévin Moneret, at iBV for their continuous support and invaluable assistance. We thank Agnès Loubat from the cytometry facility at iBV for her advice on single-cell dissociation for scRNA-Seq. We also acknowledge Matthieu Jung from the GenomEast platform, Institut de Génétique et de Biologie Moléculaire et Cellulaire (IGBMC), Illkirch, France, for his invaluable assistance with scRNA-Seq data analysis and interpretation. We further thank Antoine Fortuné and Luc Martin from the bioinformatics facility at iBV for their assistance during the initial stages of scRNA-Seq data analysis, visualization, and interpretation. Finally, we extend our special thanks to Islem Lekchiri and Gaia Bolognese for their contributions to optimizing immunostaining protocols, and to Paul Cassini for NPSC preparation.

## Funding

Fondation pour la Recherche Médicale EQU202003010222 (M.S.) Fondation de France 00123416 (M.S.)

ERA-NET NEURON ANR-21-NEU2-0003-03 (M.S.) PHENOMIN Program (M.S.)

ATIP-Avenir Program (G.G.) CNRS Starting Grant 2025 (B.Z.)

France 2030 ANR-15-IDEX-01 (B.Z.)

« Région PACA Emplois Jeunes Doctorants 2021 » (P.P.) Fondation de France 00154573 (P.P.).

Fondation Maladies Rares GenOmics Omics2024-15140 (M.B.) National Institutes of Health grant NIH R01EY036173 (S.B.)

National Institute for Health and Care Research UK NIHR301696 (P.Y.-W.-M.) LifeArc 10748 (P.Y.-W.-M.)

## Author contributions

Conceptualization: P.P., M.B., M.S. Methodology: P.P., C.B., R.M., B.Z., M.B.

Investigation: P.P., R.M., B.Z., M.B.

Data acquisition: P.P., C.B., R.M., B.Z., A.M., A.C., M.B.

Visualization: P.P., C.B., R.M., B.Z., M.B.

Funding acquisition: B.Z., P.Y.-W.-M., G.G., M.B., M.S. Supervision: P.Y.-W.-M., M.B., M.S.

Writing - original draft: M.B., M.S., P.P.

Writing - review & editing: P.P., C.B., B.Z., G.G., C.P.S., R.M., P.Y.-W.-M., S.B., M.B., M.S.

