## Supplementary data for "An NR2F1-dependent retinoic acid network controls retinal specialization in mice and reveals foveal hypoplasia in patients with BBSOAS"

Paolo Piovani *et al.*

**This PDF file includes:**

Figs. S1 to S9
Tables S1 to S5

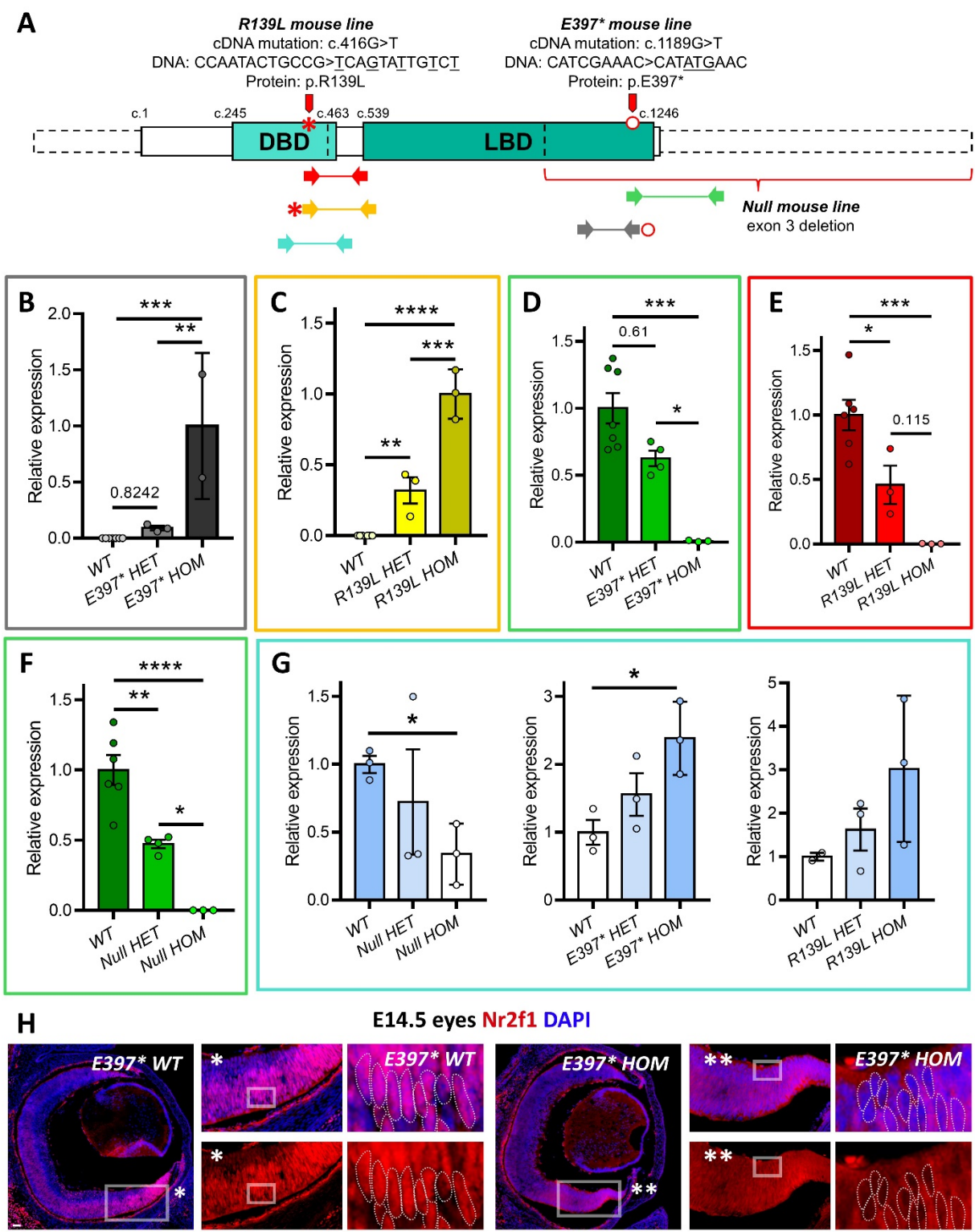

**Fig. S1. Analysis of *Nr2f1* mRNA expression and mutant protein localization.** **A)** *Nr2f1* sequence showing the genetic modifications introduced in the three mouse lines and their localization, together with the positions of the primer pairs used to detect *WT* or mutant *Nr2f1* mRNA. The linear protein structure is shown by solid lines superimposed on dotted lines indicating the corresponding mRNA exons. **B–G)** Real-time qRT-PCR quantification of *WT* or mutant *Nr2f1* mRNA expression in embryonic brains from different genotypes, using the same color code as in A to indicate the corresponding primer pairs. Primers specific for the mutant allele detected mutant *Nr2f1* expression in *E397\** and *R139L HET* and *HOM* embryos (B,C), while *WT*-specific primers revealed a corresponding reduction in *WT Nr2f1* expression (D,E). Primers targeting exon 3 detected reduced *Nr2f1* expression in *Null HET* and *HOM* embryos (F), whereas universal control primers detected increased total *Nr2f1* mRNA levels in *E397\** and *R139L* mutant embryos (G). **H)** *Nr2f1* (red) immunostaining in *E397\** *WT* and *HOM* eyes highlights the cytoplasmic localization of the truncated *Nr2f1* protein in the ventral retina. Insets show higher-magnification views highlighting the nuclear localization of *Nr2f1* in *E397\** *WT* eyes (insets marked with \*) and its cytoplasmic localization in *E397\** *HOM* eyes (insets marked with \*\*). Dashed lines in the insets delineate the position of representative RPC nuclei. Scale bars: 50 μm. Each point in graphs B-G corresponds to average expression level in one embryonic brain; n = 2-7 embryos per genotype. Statistical significance was determined by one-way ANOVA with Tukey's correction; \* p < 0.05; \*\* p < 0.01; \*\*\* p < 0.001; \*\*\*\* p < 0.0001.

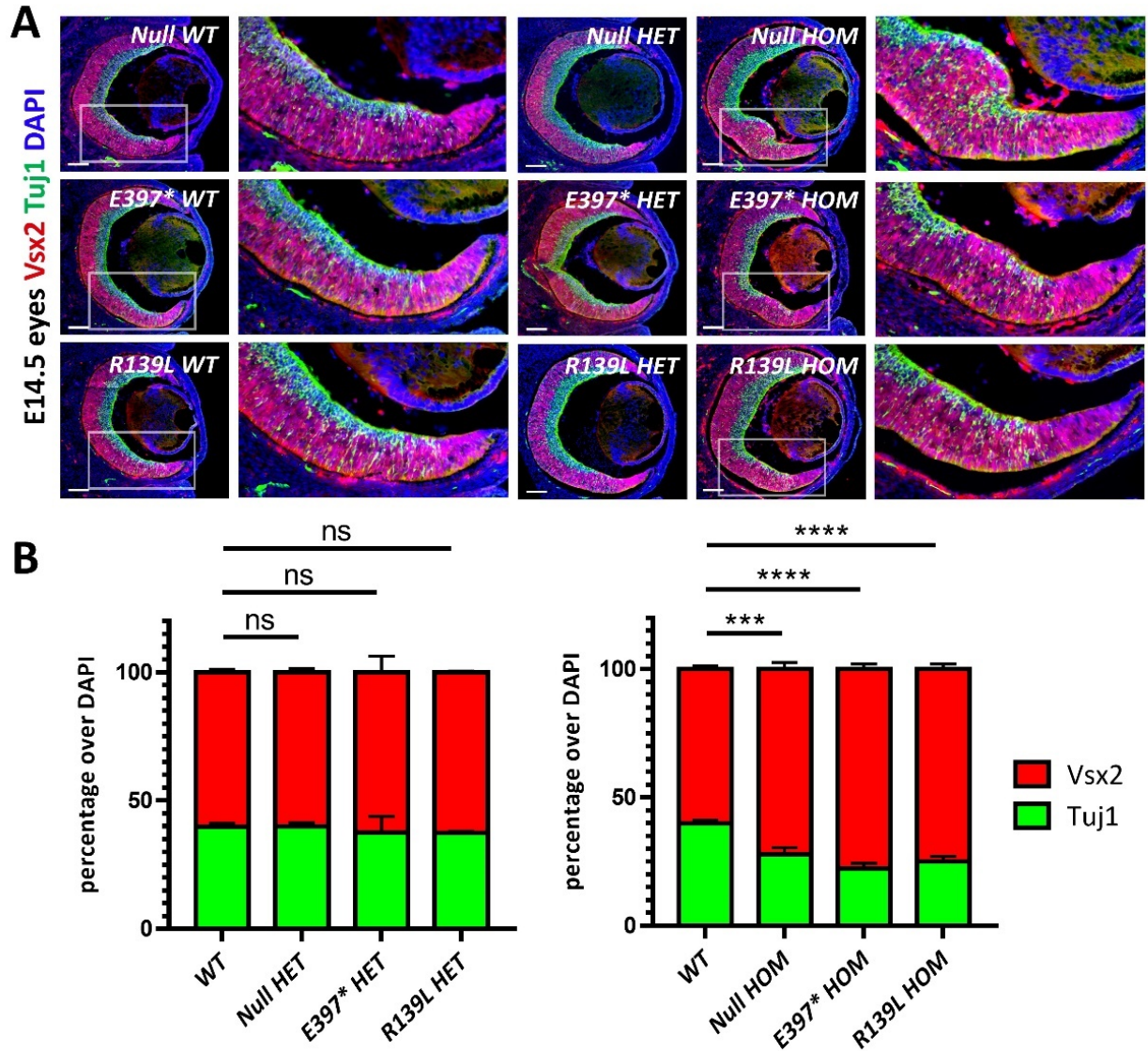

**Fig. S2. *Nr2f1* loss or mutation disrupts the balance between retinal progenitors and post-mitotic neurons.** **A)** Double immunostaining for Vsx2 (red; retinal progenitor cells) and Tuj1 (green; post-mitotic cells) in E14.5 retinæ of the indicated genotypes. **B)** Quantification of the ratio of Vsx2<sup>+</sup> retinal progenitor cells to Tuj1<sup>+</sup> differentiated post-mitotic cells in the ventral retina of *WT* and mutant embryos, revealing a significant shift toward progenitor cells in *Null*, *E397\** and *R139L HOM* mutants, consistent with delayed neurogenesis. Scale bars: 100 μm. For Graphs in B, n = 3-9 sections from n = 3-9 eyes per genotype. Statistical significance was determined by two-way ANOVA with Tukey's correction; \*\*\* p < 0.001; \*\*\*\* p < 0.0001.

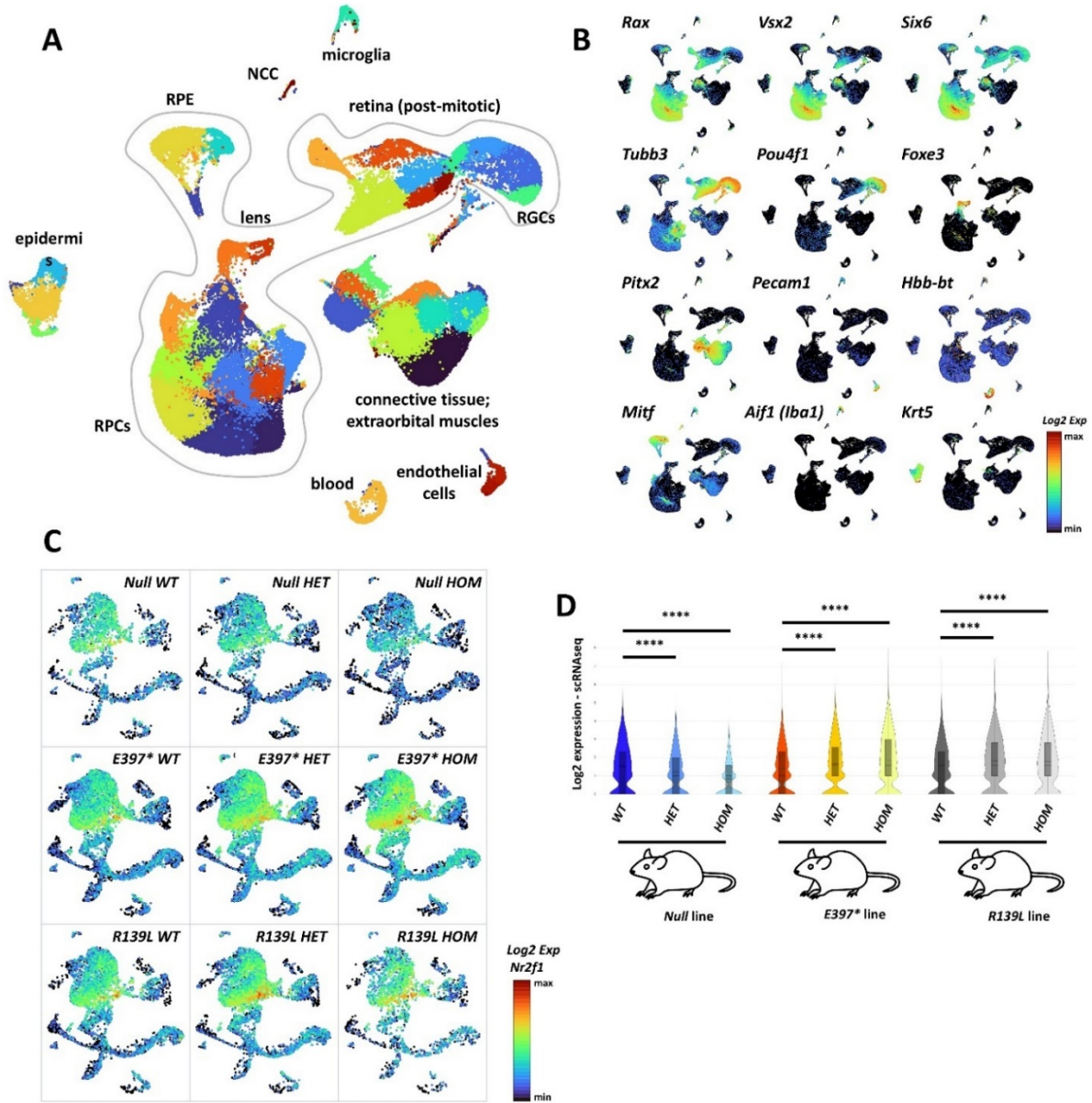

**Fig. S3. Cell-type annotation and *Nr2f1* expression in E14.5 mouse retinæ.** A,B) UMAP projection of scRNA-seq data from E14.5 mouse eyes (A), showing distinct cell populations identified by *bona fide* tissue- and cell-type-specific markers (B). Retinal progenitor cells (RPCs) were identified by *Rax*<sup>+</sup>, *Vsx2*<sup>+</sup>, and *Six6*<sup>+</sup>, whereas post-mitotic retinal populations, including retinal ganglion cells (RGCs), expressed *Tubb3* and *Pou4f1*. Retinal pigmented epithelium (RPE) was identified by *Foxe3* and *Mitf* expression. Additional non-retinal cell types arising from surrounding ocular tissues included *Krt5*<sup>+</sup> epidermal cells, *Aif1*<sup>+</sup> microglia, and *Pitx2*<sup>+</sup> connective tissue cells. C,D) *Nr2f1* mRNA expression levels in the re-clustered UMAP projection of E14.5 eye cells from the indicated genotypes (C), quantified using Loupe Browser (10x Genomics) and displayed as violin plots for each genotype (D). Statistical significance: \*\*\*\* $p < 0.0001$.

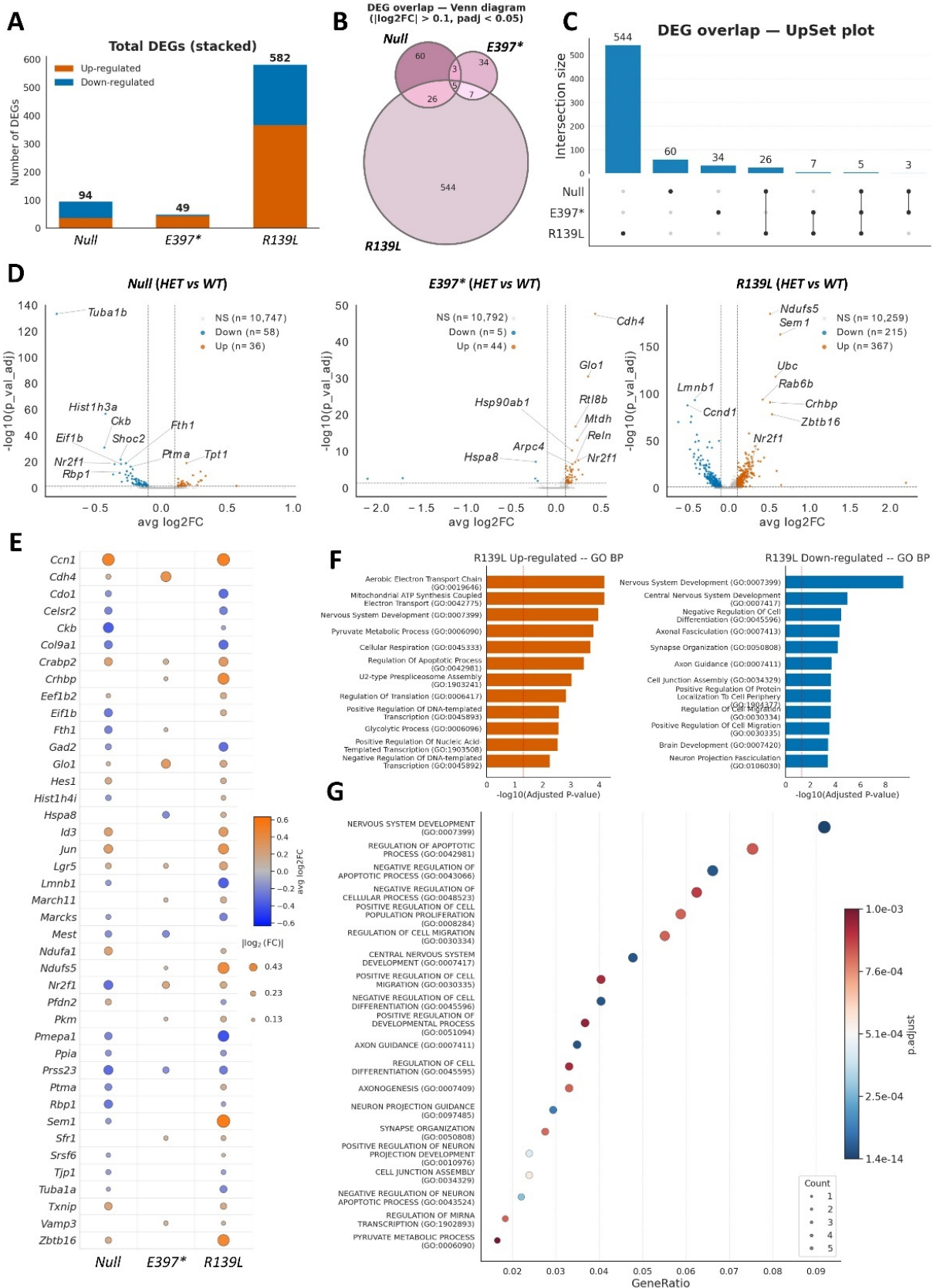

**Fig. S4. Mutation-dependent transcriptional changes in *Nr2f1* heterozygous RPCs.** **A)** Total number of up- (orange) and downregulated (blue) DEGs identified by comparing *WT* and *HET* retinal progenitor cells (RPCs) for each mouse line (*HET* vs *WT*). **B,C)** Venn diagram and intersection scheme illustrating the overlap of DEGs among the three mouse lines. Only a small number of genes are shared across all three models, whereas the *R139L* line displays several hundred DEGs. **D)** Volcano plots showing the magnitude of differential expression ($\log_2$ fold change; $\log_2FC$) and statistical significance ($-\log_{10}$ adjusted p value) for *WT* vs *HET* DEGs specific to each mouse line, as indicated. **E)** DEGs shared across two or three mouse lines, with upregulated and downregulated genes indicated in orange and blue, respectively. **F,G)** Gene ontology (GO) analysis of *R139L* DEGs, performed separately for up- and downregulated genes (F) or for the complete DEG set (G). A complete list of GO analyses for the other *WT* vs *HET* DEG subsets is provided as supplementary material.

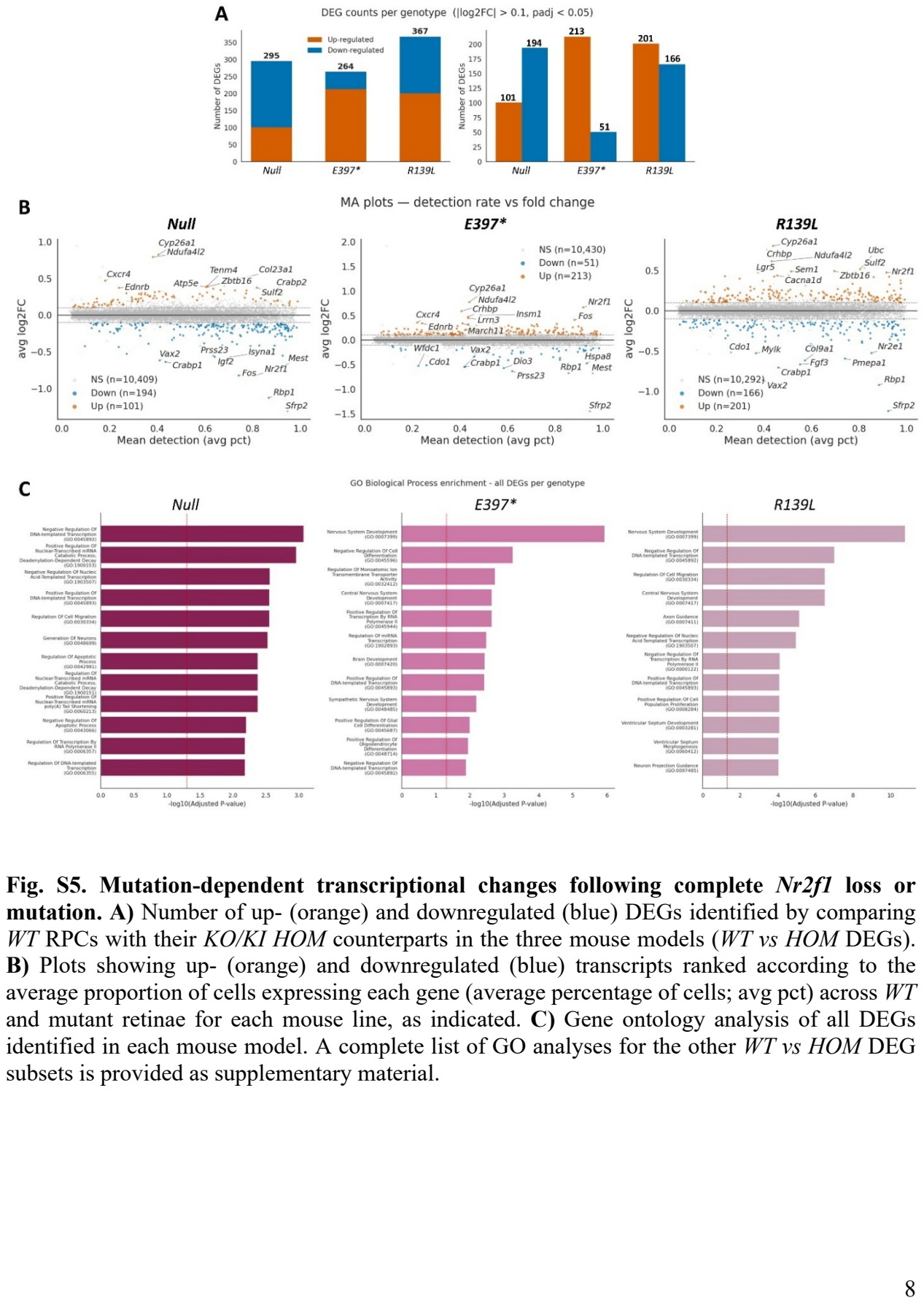

Fig. S5. Mutation-dependent transcriptional changes following complete Nr2f1 loss or mutation. A) Number of up- (orange) and downregulated (blue) DEGs identified by comparing WT RPCs with their KO/KI HOM counterparts in the three mouse models (WT vs HOM DEGs). B) Plots showing up- (orange) and downregulated (blue) transcripts ranked according to the average proportion of cells expressing each gene (average percentage of cells; avg pct) across WT and mutant retinae for each mouse line, as indicated. C) Gene ontology analysis of all DEGs identified in each mouse model. A complete list of GO analyses for the other WT vs HOM DEG subsets is provided as supplementary material.

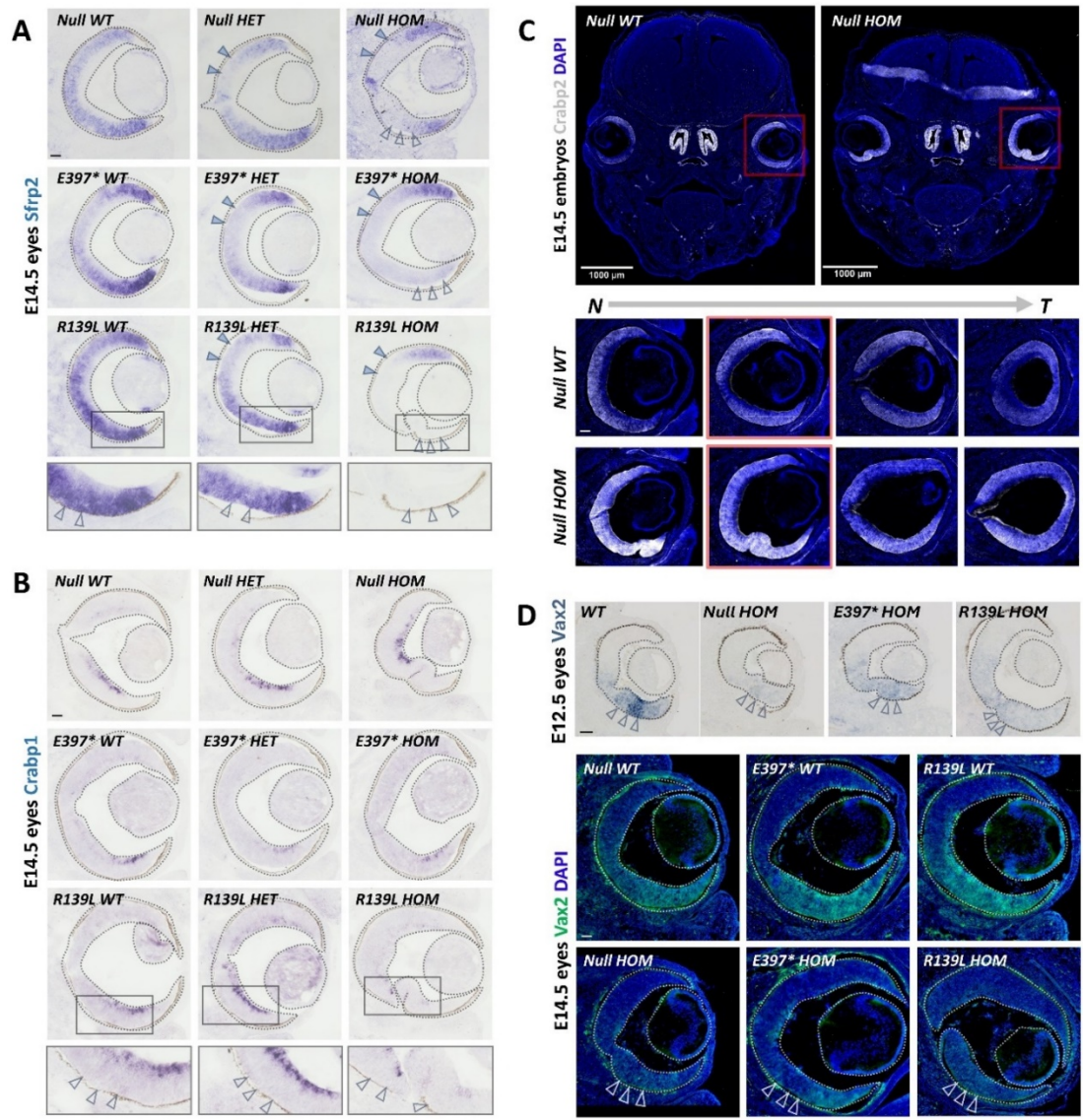

**Fig. S6. Altered spatial expression of *Nr2f1*-dependent regional patterning genes in mutant retinae.** **A)** *In situ* hybridization for *Sfrp2* transcript in E14.5 mouse retinae of the indicated genotypes. Blue arrowheads indicate reduced dorsal *Sfrp2* expression, already detectable in *HET* embryos, whereas white arrowheads indicate the ventral-most retina, where *Sfrp2* expression was completely lost specifically in *R139L HOM* mutants. **B)** *In situ* hybridization for *Crabp1* in E14.5 mouse retinae of the indicated genotypes. White arrowheads indicate reduced *Crabp1* expression in RPCs of *R139L* mutants, with a concomitant reduction in RGCs. **C)** Immunostaining for *Crabp2* (white) in E14.5 embryo heads, showing increased *Crabp2* expression in the ventral retina of *Null HOM* mutants at different positions along the naso (N)–temporal (T) axis. **D)** *In situ* hybridization for *Vax2* in E12.5 retinae and *Vax2* immunostaining (green) in E14.5 retinae of the indicated genotypes, showing decreased expression in the ventral retina (arrowheads) of mutant embryos. Scale bars: 100 μm, except upper panel in C, 1000 μm.

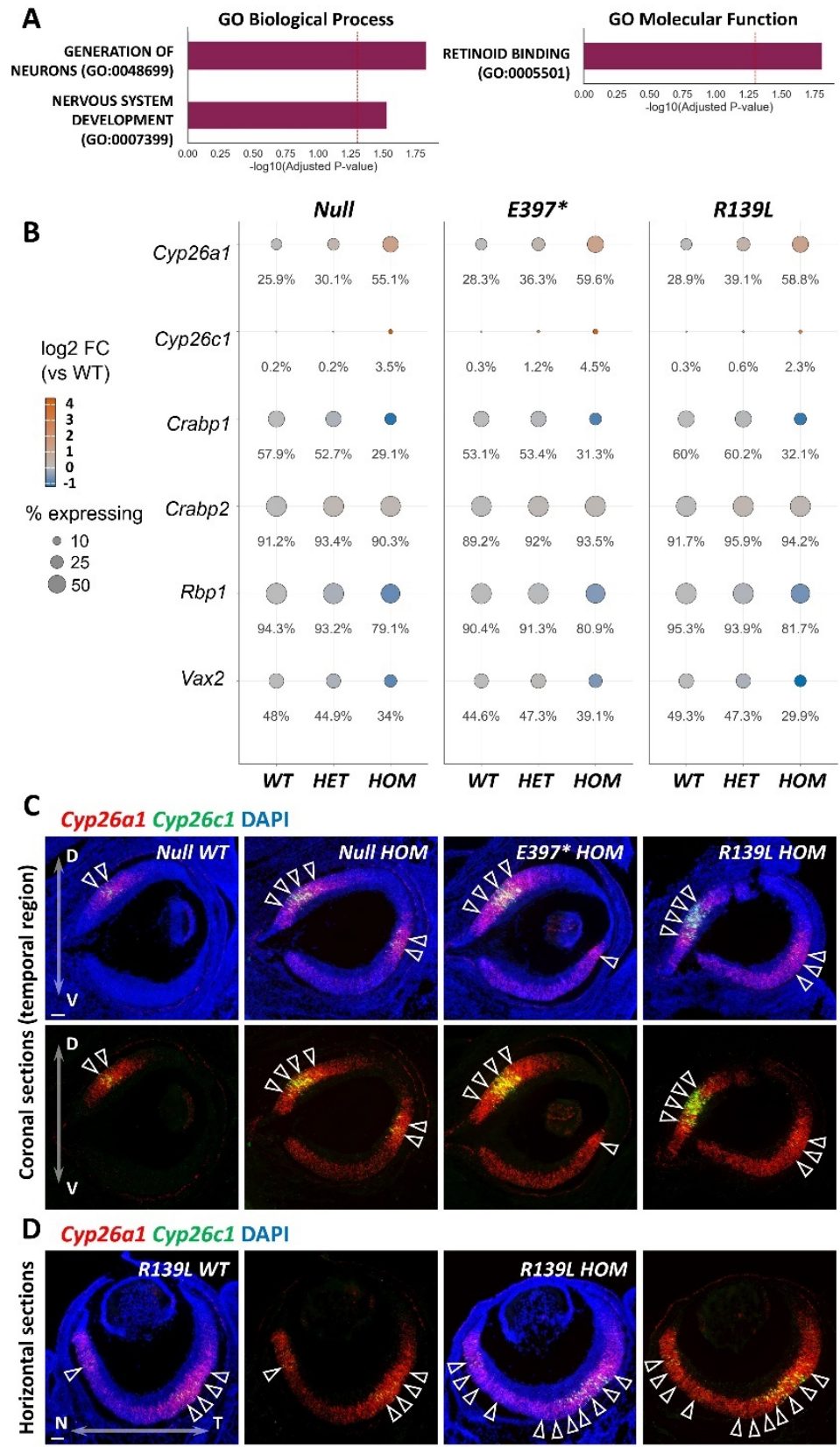

**Fig. S7. Gene ontology and spatial analysis of *Nr2f1*-dependent RA pathway dysregulation.** **A)** Gene ontology analysis of the 67 shared DEGs, highlighting “generation of neurons” and “nervous system development” as enriched GO biological processes, and “retinoid binding” as a significantly enriched molecular function. **B)** Dot plot showing the proportion of cells expressing the RA-related genes *Cyp26a1*, *Cyp26c1*, *Crabp1*, *Crabp2*, *Rbp1*, and *Vax2*, together with their fold change in *WT*, *HET* and *HOM KO/KI* animals for the indicated genotypes. **C)** RNAscope *in situ* hybridization for *Cyp26a1* (red) and *Cyp26c1* (green) in temporal-most coronal sections of E14.5 retinae from the indicated genotypes. Arrowheads indicate equato-dorsal retinal regions and ventral CMZ areas. Upregulation of both paralogs, together with ectopic *Cyp26a1* expression, is particularly pronounced in the ventral retina (D = dorsal, V = ventral). **D)** RNAscope *in situ* hybridization for *Cyp26a1* (red) and *Cyp26c1* (green) in horizontal sections of E14.5 *WT* and *R139L HOM KI* retinae. Arrowheads point to high *Cyp26a1* expression. Ectopic upregulation of *Cyp26a1* is detected in the nasal retina (N = nasal, T = temporal). Scale bars: 50 μm.

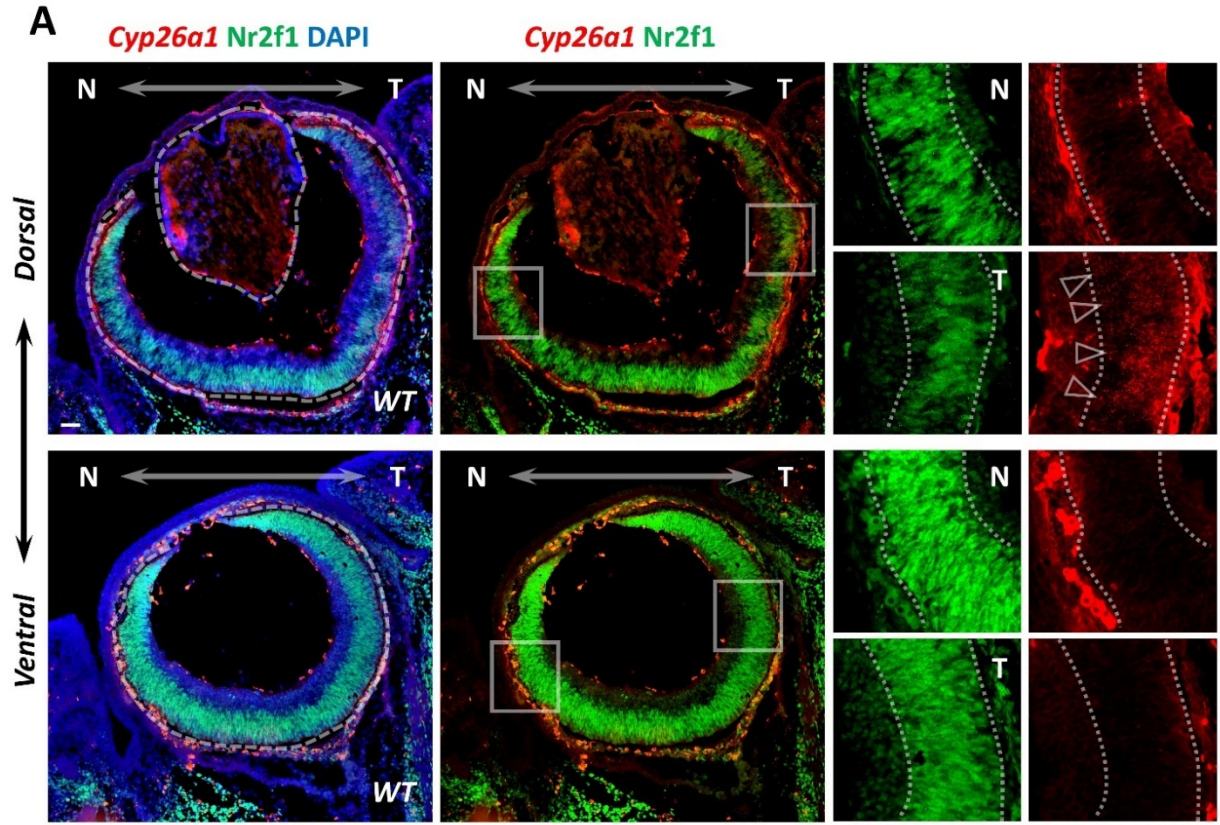

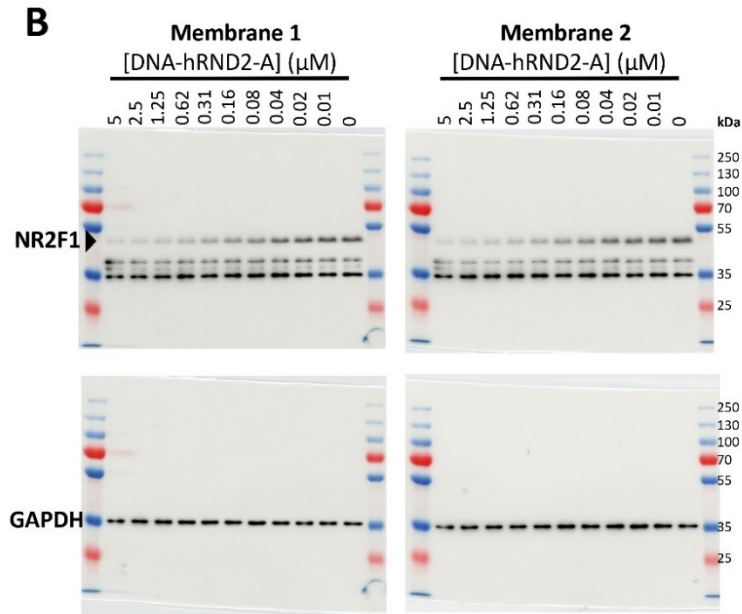

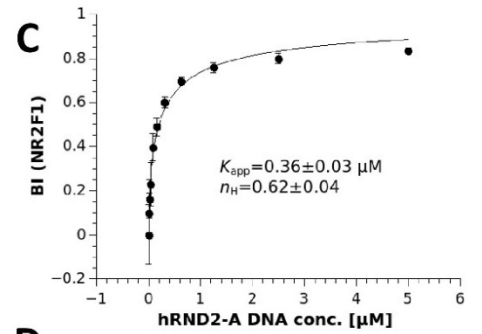

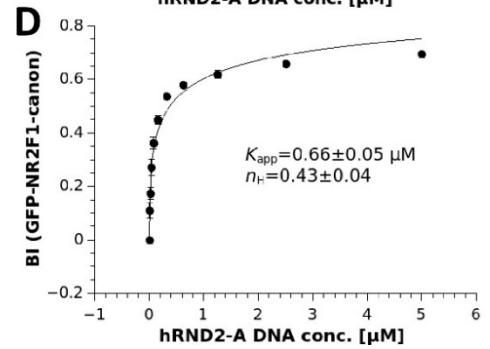

**Fig. S8. Complementary N-T expression gradients of Nr2f1 and *Cyp26a1* and validation of NR2F1 DNA-binding activity by native holdup.** **A)** RNAscope *in situ* hybridization for *Cyp26a1* (red) combined with Nr2f1 immunostaining (green) on E14.5 WT eye horizontal sections at the indicated dorsal and ventral levels. Nr2f1 exhibits a nasal (N)-to-temporal (T) decreasing gradient, more pronounced at the dorsal level. Conversely, sparse *Cyp26a1* expression (arrowheads in inset) is detected predominantly in the temporal retina, where Nr2f1 expression levels are lower. **B–D)** Native holdup analysis using endogenous NR2F1 from neural progenitor cells and GFP-tagged NR2F1 expressed in HEK293T cells. An NR2F1 recognition sequence from the human *RND2* gene was used as DNA bait. NR2F1 prey proteins were captured directly from cell extracts and detected by Western blotting or GFP fluorescence. In B, Western blot analysis of native holdup performed with endogenous NR2F1 from neural progenitor stem cell (NPSC) extracts (day 12 of differentiation). C shows the binding curve derived from the Western blot quantification shown in B (n = 2). In D, the binding curve obtained from native holdup using GFP-NR2F1 expressed in HEK293T cells (n = 3). BI, bound fraction intensity; Kapp, apparent affinity; nH, Hill coefficient. Scale bars: 50 μm.

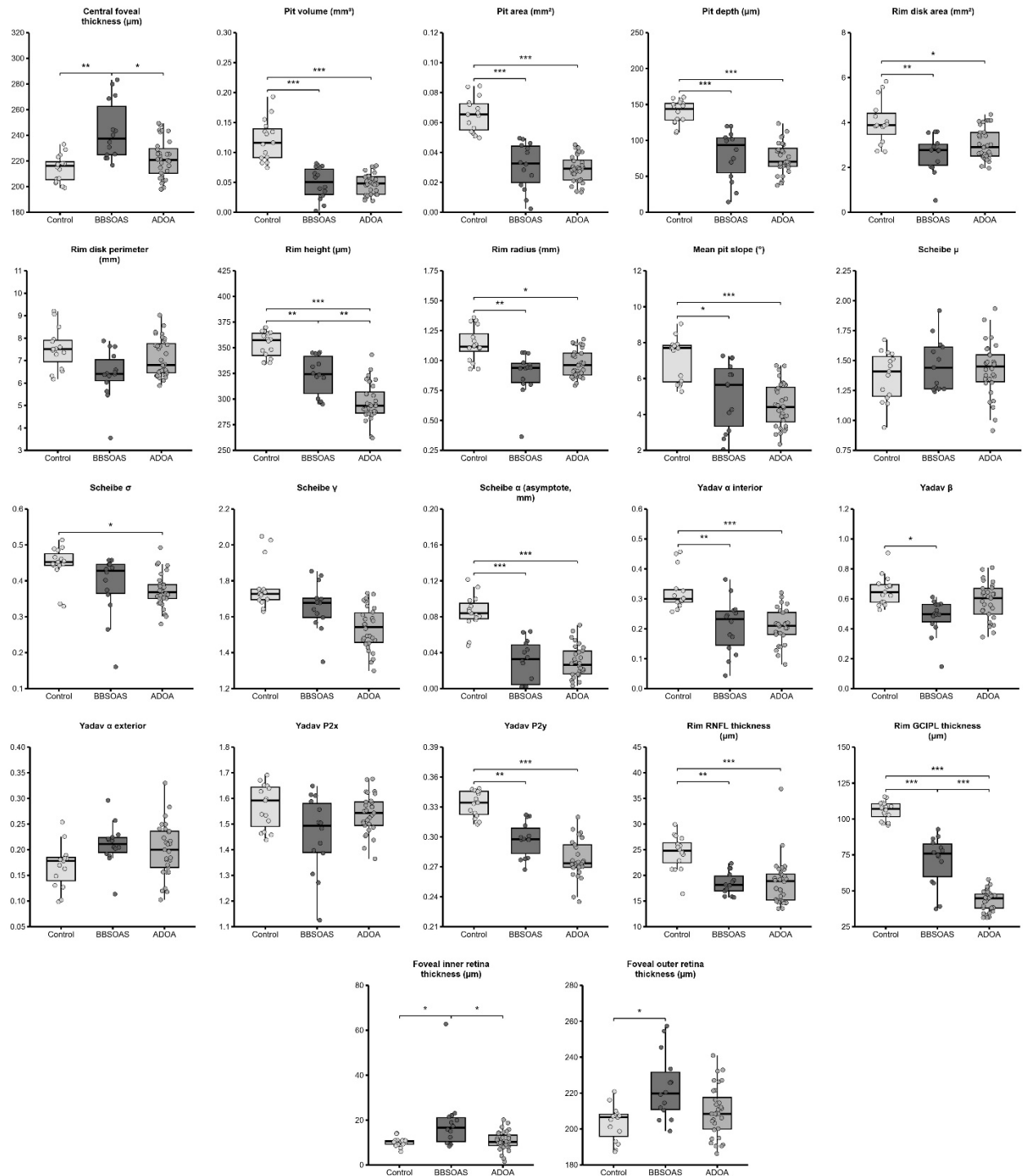

**Fig. S9. Comparison of foveal morphology across BBSOAS, ADOA, and healthy controls.** Boxplots comparing 22 quantitative foveal pit morphology parameters among individuals with BBSOAS (n = 14 eyes from 7 patients; dark grey), autosomal dominant optic atrophy (ADOA; n = 32 eyes from 16 patients; grey), and age-matched healthy controls (n = 16 eyes from 8 individuals; light grey). Boxes represent the interquartile range, with the median indicated. Statistical significance was assessed using linear mixed-effects models accounting for measurements from both eyes, with P values adjusted for multiple comparisons using the Benjamini–Hochberg procedure (\*P < 0.05, \*\*P < 0.01, \*\*\*P < 0.001). Parameters were grouped into four categories: (i) foveal pit morphology derived from radial retinal thickness profiles (central foveal thickness, pit volume/area/depth, rim disk area/perimeter, rim height/radius, mean pit slope); (ii) Scheibe et al. radial parametric fit coefficients ($\mu$, $\sigma$, $\gamma$, $\alpha$); (iii) Yadav et al. cubic-Bézier fit coefficients ($\alpha$ interior/exterior, $\beta$, P2x, P2y); and (iv) retinal layer thicknesses measured at the pit rim (RNFL, GCIPL) and foveal center (inner retina = ILM to IPL/INL; outer retina = IPL/INL to BM). Demographic characteristics of patients with BBSOAS, patients with ADOA, and control individuals included in the analysis are provided in *Table S1*; *NR2F1* and *OPA1* pathogenic variants in the patients included in this analysis are listed in *Tables S2* and *S4*, respectively. Summary statistics for the foveal parameters shown in the graphs are provided in *Table S5*.

| Cohort | n_patients | n_eyes | n_female | pct_female | age_mean | age_sd | age_range | summary |
| --- | --- | --- | --- | --- | --- | --- | --- | --- |
| Control | 8 | 16 | 4 | 50 | 281 | 5 | 23-37 | n=8 (16 eyes), 50% female, age at scan 28.1 +/- 5.0 (23-37) |
| ADOA | 16 | 32 | 8 | 50 | 403 | 94 | 23-57 | n=16 (32 eyes), 50% female, age at scan 40.3 +/- 9.4 (23-57) |
| BBSOAS | 7 | 14 | 4 | 57 | 361 | 209 | 16-75 | n=7 (14 eyes), 57% female, age at scan 36.1 +/- 20.9 (16-75) |

**Table S1. Demographic summary of control, ADOA and BBSOAS individuals.**

BBSOAS\_patient\_genotype\_list

| Study ID | Gene | cDNA (HGVS) | Protein | Family | In original paper |
| --- | --- | --- | --- | --- | --- |
| **BB-01** | NR2F1 | c.1115T>C | p.(Leu372Pro) | F1 | Yes |
| **BB-02** | NR2F1 | c.1115T>C | p.(Leu372Pro) | F1 | Yes |
| **BB-03** | NR2F1 | c.1115T>C | p.(Leu372Pro) | F1 | Yes |
| **BB-04** | NR2F1 | c.1118_1123del | p.(Arg373_Leu374del) | F2 | Yes |
| **BB-05** | NR2F1 | c.1036_1047del | p.(Glu346_Gln349del) | F3 | Yes |
| **BB-06** | NR2F1 | c.237G>C | p.(Gln79His) | F4 | No |
| **BB-07** | NR2F1 | (not recorded) | (not recorded) | F5 | No |

**Table S2. *NR2F1* pathogenic variants in individuals with BBSOAS included in the foveal morphometric analysis**

BBSOAS\_vs\_Control\_summary\_table

| Metric | Control (8 patients, 16 eyes) | BBSOAS (7 patients, 14 eyes) | p (BH-adj) |
| --- | --- | --- | --- |
| Central foveal thickness ($\mu\text{m}$) | 214.16 ± 10.33 | 243.37 ± 22.98 | 0.015 |
| Foveal inner retina thickness ($\mu\text{m}$) | 10.23 ± 2.10 | 18.94 ± 13.62 | 0.065 |
| Foveal outer retina thickness ($\mu\text{m}$) | 203.32 ± 9.86 | 223.42 ± 18.41 | 0.025 |
| Mean pit slope ($^{\circ}$) | 7.12 ± 1.20 | 5.07 ± 1.85 | 0.039 |
| Pit area ($\text{mm}^2$) | 0.07 ± 0.01 | 0.03 ± 0.02 | 0.002 |
| Pit depth ($\mu\text{m}$) | 139.18 ± 16.11 | 79.94 ± 34.64 | 0.003 |
| Pit volume ($\text{mm}^3$) | 0.12 ± 0.03 | 0.05 ± 0.03 | 0.003 |
| Rim GCiPL thickness ($\mu\text{m}$) | 105.78 ± 6.40 | 70.91 ± 17.41 | 0.002 |
| Rim RNFL thickness ($\mu\text{m}$) | 24.46 ± 3.33 | 18.44 ± 2.19 | 0.002 |
| Rim disk area ($\text{mm}^2$) | 4.03 ± 0.97 | 2.60 ± 0.83 | 0.015 |
| Rim disk perimeter (mm) | 7.53 ± 0.91 | 6.40 ± 1.10 | 0.044 |
| Rim height ($\mu\text{m}$) | 353.34 ± 12.64 | 323.31 ± 19.07 | 0.010 |
| Rim radius (mm) | 1.14 ± 0.14 | 0.89 ± 0.18 | 0.015 |
| Scheibe $\alpha$ (asymptote, mm) | 0.08 ± 0.02 | 0.03 ± 0.02 | 0.002 |
| Scheibe $\gamma$ | 1.77 ± 0.13 | 2.80 ± 4.27 | 0.358 |
| Scheibe $\mu$ | 1.37 ± 0.21 | 2.57 ± 4.15 | 0.308 |
| Scheibe $\sigma$ | 0.45 ± 0.05 | 0.39 ± 0.09 | 0.150 |
| Yadav P2x | 1.57 ± 0.09 | 1.46 ± 0.15 | 0.069 |
| Yadav P2y | 0.33 ± 0.01 | 0.30 ± 0.02 | 0.003 |
| Yadav $\alpha$ exterior | 0.17 ± 0.04 | 0.21 ± 0.04 | 0.044 |
| Yadav $\alpha$ interior | 0.32 ± 0.07 | 0.21 ± 0.09 | 0.015 |
| Yadav $\beta$ | 0.66 ± 0.10 | 0.48 ± 0.12 | 0.014 |

**Table S3. Summary table of the foveal morphological parameters measured for both control and individuals with BBSOAS.**

ADOA\_patient\_genotype\_list

| Study ID | Gene | cDNA (HGVS) | Family |
| --- | --- | --- | --- |
| **ADOA-01** | OPA1 | c.1241C>G | F1 |
| **ADOA-02** | OPA1 | c.1241C>G | F1 |
| **ADOA-03** | OPA1 | c.1149G>A |  |
| **ADOA-04** | OPA1 | c.2708_2711del |  |
| **ADOA-05** | OPA1 | c.869G>A |  |
| **ADOA-06** | OPA1 | c.2818+1G>A |  |
| **ADOA-07** | OPA1 | c.2521-2A>G |  |
| **ADOA-08** | OPA1 | c.2873_2876delTTAG |  |
| **ADOA-09** | OPA1 | c.2706_2711del |  |
| **ADOA-10** | OPA1 | c.1140G>A |  |
| **ADOA-11** | OPA1 | c.870+2T>A |  |
| **ADOA-12** | OPA1 | Deletion of exons 2–4 (~2.6 kb) |  |
| **ADOA-13** | OPA1 | c.1212+1G>A |  |
| **ADOA-14** | OPA1 | c.1491_1493delins37 |  |
| **ADOA-15** | OPA1 | c.899G>A |  |
| **ADOA-16** | OPA1 | c.1187T>G |  |

**Table S4. *OPA1* pathogenic variants in individuals with ADOA included in the foveal morphometric analysis.**

Control\_BBSOAS\_ADOA\_summary\_table

| Metric | Control (8 patients, 16 eyes) | BBSOAS (7 patients, 14 eyes) | ADOA (16 patients, 32 eyes) | Omnibus p (BH-adj) | BBSOAS vs Control (Tukey) | ADOA vs Control (Tukey) | BBSOAS vs ADOA (Tukey) |
| --- | --- | --- | --- | --- | --- | --- | --- |
| Central foveal thickness (μm) | 214.16 ± 10.33 | 243.37 ± 22.98 | 221.56 ± 15.03 | 0.008 | 0.005 | 0.554 | 0.017 |
| Pit volume (mm<sup>3</sup>) | 0.12 ± 0.03 | 0.05 ± 0.03 | 0.05 ± 0.02 | <0.001 | <0.001 | <0.001 | 0.997 |
| Pit area (mm<sup>2</sup>) | 0.07 ± 0.01 | 0.03 ± 0.02 | 0.03 ± 0.01 | <0.001 | <0.001 | <0.001 | 0.933 |
| Pit depth (μm) | 139.18 ± 16.11 | 79.94 ± 34.64 | 75.66 ± 21.18 | <0.001 | <0.001 | <0.001 | 0.919 |
| Rim disk area (mm<sup>2</sup>) | 4.03 ± 0.97 | 2.60 ± 0.83 | 3.04 ± 0.70 | 0.005 | 0.003 | 0.012 | 0.453 |
| Rim disk perimeter (mm) | 7.53 ± 0.91 | 6.40 ± 1.10 | 7.13 ± 0.84 | 0.056 | 0.037 | 0.472 | 0.173 |
| Rim height (μm) | 353.34 ± 12.64 | 323.31 ± 19.07 | 297.22 ± 18.03 | <0.001 | 0.007 | <0.001 | 0.007 |
| Rim radius (mm) | 1.14 ± 0.14 | 0.89 ± 0.18 | 0.98 ± 0.11 | 0.006 | 0.003 | 0.022 | 0.353 |
| Mean pit slope (°) | 7.12 ± 1.20 | 5.07 ± 1.85 | 4.52 ± 1.20 | 0.002 | 0.022 | <0.001 | 0.662 |
| Schelbe μ | 1.37 ± 0.21 | 2.57 ± 4.15 | 1.43 ± 0.23 | 0.198 | 0.273 | 0.995 | 0.217 |
| Schelbe σ | 0.45 ± 0.05 | 0.39 ± 0.09 | 0.37 ± 0.05 | 0.036 | 0.162 | 0.022 | 0.826 |
| Schelbe γ | 1.77 ± 0.13 | 2.80 ± 4.27 | 1.54 ± 0.12 | 0.178 | 0.384 | 0.932 | 0.160 |
| Schelbe α (asymptote, mm) | 0.08 ± 0.02 | 0.03 ± 0.02 | 0.03 ± 0.02 | <0.001 | <0.001 | <0.001 | 0.999 |
| Yadav α interior | 0.32 ± 0.07 | 0.21 ± 0.09 | 0.21 ± 0.06 | 0.001 | 0.004 | <0.001 | 0.997 |
| Yadav β | 0.66 ± 0.10 | 0.48 ± 0.12 | 0.58 ± 0.12 | 0.022 | 0.011 | 0.308 | 0.107 |
| Yadav α exterior | 0.17 ± 0.04 | 0.21 ± 0.04 | 0.20 ± 0.05 | 0.112 | 0.112 | 0.181 | 0.820 |
| Yadav P2x | 1.57 ± 0.09 | 1.46 ± 0.15 | 1.54 ± 0.08 | 0.056 | 0.046 | 0.682 | 0.120 |
| Yadav P2y | 0.33 ± 0.01 | 0.30 ± 0.02 | 0.28 ± 0.02 | <0.001 | 0.002 | <0.001 | 0.057 |
| Rim RNFL thickness (μm) | 24.46 ± 3.33 | 18.44 ± 2.19 | 18.71 ± 4.43 | <0.001 | 0.002 | <0.001 | 0.978 |
| Rim GCIPL thickness (μm) | 105.78 ± 6.40 | 70.91 ± 17.41 | 42.95 ± 7.18 | <0.001 | <0.001 | <0.001 | <0.001 |
| Foveal inner retina thickness (μm) | 10.23 ± 2.10 | 18.94 ± 13.62 | 10.68 ± 4.19 | 0.019 | 0.028 | 0.986 | 0.015 |
| Foveal outer retina thickness (μm) | 203.32 ± 9.86 | 223.42 ± 18.41 | 209.68 ± 13.87 | 0.033 | 0.020 | 0.477 | 0.097 |

**Table S5. Summary table of the foveal morphological parameters measured for control, ADOA and BBSOAS individuals.**
